# Diversity and abundance of tree microhabitats in the tropical forests of southern Western Ghats, India

**DOI:** 10.1101/2024.03.23.586393

**Authors:** Bharati Patel, Sreejith Sivaraman, T.K. Hrideek, Peroth Balakrishnan

## Abstract

Tree microhabitats (TMHs) are proven tools for assessing and monitoring diversity. These structures on trees are potential indicators of biota, but there is a huge gap in TMH-related knowledge from the tropical regions, the cradles of biodiversity. Thus, an inventory was made to document the TMHs in the tropical forests of southern Western Ghats, India. In evergreen forests, 3,637 TMH host and 450 cavity host trees were identified from the 6,363 trees sampled. The density of TMHs was 972.57±341.25 ha^-1^ and cavity density was 63.13±11.91 ha^-1^. In moist deciduous forests, out of 1,545 trees sampled, 1,108 hosted TMHs and 212 trees hosted cavities. The density of TMHs was 493.67±133.28 ha^-1^ and cavity density was 42.00±15.01 ha^-1^. TMHs were categorised into nine categories and 33 sub-categories. The TMH and cavity occurrences were significantly influenced by species richness, stand density, basal area, diameter and height of trees, and density of healthy, unhealthy and dead trees. Dominant and codominant individuals hosted more TMHs and cavities in the deciduous stands while in evergreen, intermediate and overtopped trees had more TMHs and intermediate and codominant had more cavities. In both the habitats the important species of the habitat were also major hosts for TMHs and cavities.

**Highlights:**

- Primary inventory of diversity of TMHs and their host trees in tropical forests
- TMH and cavity occurrences are significantly influenced by stand characteristics
- Key species in the habitats also form major hosts for TMHs and cavities
- Dominant and codominant individuals host more TMHs and cavities in deciduous stands
- Intermediate and overtopped trees host more TMHs, intermediate and codominant host more cavities in evergreen

## 1. Introduction

Tropical rainforests are known for their unparalleled complexity and species diversity due to their diverse abiotic factors, therefore making them a hub of biological research for understanding natural processes (Gentry, 1992). Biotic and abiotic elements jointly shape the horizontal and vertical structures of these forests (Martinez-Yrizar et al., 2000; Tews et al., 2004), with the spatial patterns, size distribution, and height variations in trees greatly influencing structural heterogeneity (Sabatini et al., 2015). This structural diversity fosters numerous habitats, supporting a multitude of organisms (Rapp, 2003) and validating the “habitat heterogeneity hypothesis” for tropical forests.

The immense ecological complexity of these habitats, evolved over millions of years, poses challenges in prioritizing conservation areas, assessing management efforts, and gauging restoration progress. Integrating the concept of indicator species is one of the keys to such issues in conservation and management programmes. Given the constraints associated with species identification and taxon-based indicators, there’s a call for a focused, evidence-based approach (Gardner et al., 2008). Structure-based biodiversity indicators, including forest connectivity, stand heterogeneity, and ecosystem functions (Noss, 1990; Hunter, 1999; Lindenmayer et al., 2000), are proposed as alternatives. Notably, structured Pacific coastal old-growth forests illustrate this concept, harbouring elusive organisms like bats, lichens, fungi, and invertebrates (Thomas, 1988; Schowalter, 1995; McCune et al., 2000; Addison et al., 2003; Smith et al., 2003).

The structure-based indicators, especially those related to microhabitat niches on individual trees, have proven highly correlated with the abundance of forest species and ecosystem functions (Thomas et al., 1979; Harmon et al., 1986; Vuidot et al., 2011; Parsons et al., 2003; Remm et al., 2006; Bouget et al., 2013; Regnery et al., 2013; Paillet et al., 2018; Parisi et al., 2021; Asbeck et al., 2022). Quantifying and mapping these microhabitats can serve as an indicator for these hidden organisms.

Tree microhabitats (TMHs) are specific above-ground morphological features found on tree trunks and branches in forest ecosystems, often considered “keystone structures” vital for various dependent taxa (Remm and Lõhmus, 2011). They host a wide range of flora and fauna, facilitating ecosystem services such as predation and pollination (Kumar and Singh, 2010; Wordley et al., 2017).

Documenting TMH diversity and its dependent fauna provides an alternative for biodiversity monitoring, habitat prioritization, and sustainable forest management, as demonstrated in temperate, Mediterranean, and boreal forests (Michel and Winter, 2009; Regnery et al., 2013; Paillet et al., 2018; Moreira-Arce et al., 2021). This approach is proposed as a valuable tool for conserving biological diversity in both natural and managed forest stands (Schuck et al., 2005; Franklin and Johnson 2012; Augustynczik et al., 2019; Oettel and Lapin 2021; Asbeck et al., 2022; Martin et al., 2022). Given the dire threat to tropical zones due to biodiversity loss resulting from deforestation and degradation (Quesada and Stoner, 2004; Peres, 2006; Stork et al., 2009; Wilkie et al., 2011; Laurance, 2013; França et al., 2020), the application of structure-based indicators in tropical forests is of utmost ecological significance.

While extensive research has typified TMHs in temperate and Mediterranean regions (Winter and Moller, 2008; Vuidot et al., 2011; Larrieu et al., 2012; Regnery et al., 2013; Larrieu et al., 2014; Kraus et al., 2016), there’s a notable gap in understanding TMH diversity and distribution in tropical forests and their role as surrogates for dependent fauna. Some studies have explored related forest components such as epiphytes, lichens, tree cavities, and deadwoods (Stegge and Cornelissen, 1989; Annaselvam and Parthasarathy, 2001; Muñoz et al., 2003; Padmawathe et al., 2004; Rout et al., 2010; Ding et al., 2016; Wolseley and Aguirre-Hudson, 1997; Komposch and Hafellner, 2000; Kalb et al., 2009; Prajapati et al., 2013; Benítez et al., 2015; Liu et al., 2019). However, the identification and quantification of TMHs as biodiversity surrogates in tropical regions remains limited (Grove, 2002; Remm and Lõhmus, 2011).

This study aims to estimate the diversity and abundance of TMHs in the tropical forests of India, recognizing their association with stand complexity and heterogeneity. It is hypothesized that evergreen stands, characterized by higher density and species richness, will exhibit greater TMH diversity and abundance. Additionally, within evergreen stands, plots and transects with higher species diversity are expected to host more TMHs.

## 2. Materials and Methods

### 2.1. Study area

Two representative vegetation types, namely, evergreen forests and moist deciduous forests from the southern Western Ghats were selected for the inventory. Sampling was done in protected forest areas to avoid bias due to anthropogenic activities. The areas selected were Silent Valley National Park (SVNP, 11° 2’N and 11° 13’N and 76° 24’E and 76° 32’E), Muthikkulam Reserve Forest (MRF, 10°56’N and 10°59’N and 76°41’E and 76°45’ E) and Peechi Vazhani Wildlife Sanctuary (PVWLS, 10°28’N to 10°38’N and 76°18’E to 76° 28’E) in the Kerala state in southern India (Figure 1). SVNP has a core zone of 89.52 sq km with a 148 sq km buffer zone making the total area as 237.52 sq km. The elevation of the park ranges from 70 to 2,383 m asl, annual temperature ranges from 9° C to 38° C. The region receives a bulk of both south-west and north-east monsoon showers with an average rainfall >5,000 mm. These climates generously support the representative rainforests of the southern Western Ghats, the West-coast tropical evergreen forests, Southern subtropical broadleaved hill forests, Southern montane wet temperate forests and grasslands and harbour many endemic flora as well as fauna. MRF is under the Mannarkkad Forest Division and forms the southern part of the Nilgiri Biosphere Reserve. The reserve has an area of 63.86 sq km with an elevation ranging from 610 to 2,065 m and a temperature range from 10° C in winters to 30–32° C in summers. These forests receive both south–west and north–east monsoons, with an average rainfall of 2,000 to 4,500 mm. These reserve forests are dominated by West–coast tropical evergreen forests, West–coast tropical semi–evergreen forests, Southern montane wet temperate forests and grasslands. It is also identified as an important bird area with high diversity (BirdLife International, 2000). In both SVNP and MRF sampling was done in the evergreen forests. PVWLS has an extent of 125 sq km. The terrain is undulating with elevation varying from 45 to 900 m and annual temperature ranges from 12°C to 39°C. This area receives an average rainfall of 3,000 mm. The dominant vegetation is tropical moist–deciduous forests and semi– evergreen forests. Sampling was done in the moist deciduous forests. The inventories were made from December 2017 to June 2020.

**Figure 1:**
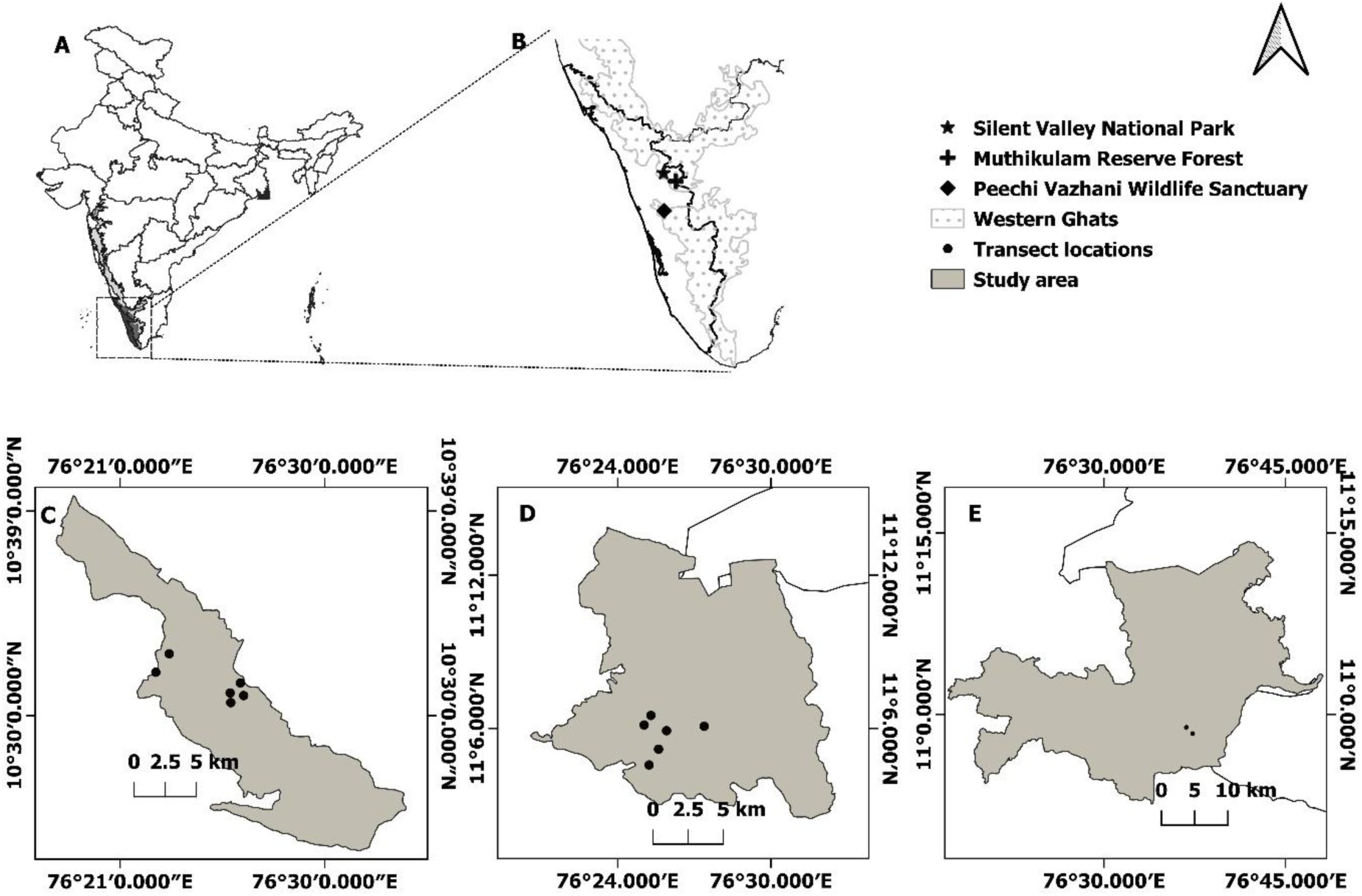
Map of study area showing locations of sampling sites: (A) India, (B) Southern Western Ghats of Kerala region, (C) Peechi-Vazhani Wildlife Sanctuary, (D) Silent Valley National Park, (E) Muthikkulam Reserve Forest.

### 2.2. Sampling design

A pilot survey was conducted in 4 ha area of Silent Valley National Park (SVNP) and rectangular plots of 50 m × 20 m were found to be most suitable for documenting the tree microhabitats (TMHs). Other researchers working on TMHs also suggested that rectangular plots work better for forest structures which are assumed to have a clustered distribution (Bate et al., 1999; Arellano et al., 2016). The study area was stratified as evergreen forests and moist deciduous forests to compare the influence of the type of habitat and stand characteristics on the diversity and density of TMHs. In both habitats sampling in 6 ha area of natural forest stands (6 transects of each 1 ha area) was carried out following the standard forest biometry procedures (Chaturvedi and Khanna, 2000; Husch et al., 2003). For sampling, transects were marked, each being 1,000 m long and 20 m wide on either side of the centreline (Figure 2). Each transect was divided into forty, 50 m × 20 m (0.1 ha) quadrats and 10 alternate quadrats from both sides of the centreline were selected for sampling (Arellano et al., 2016). Trees were surveyed for tree microhabitats in 60 selected quadrats, 6 ha each in evergreen forests of SVNP and moist deciduous forests of PVWLS, but only 2 ha in evergreen forests of MRF due to logistic reasons. In each quadrat all the trees with diameter at breast height (dbh) ≥ 0.10 m and snag or stump with height ≥ 1 m were sampled (Bate et al., 1999). In both the habitats all the live and dead trees sampled were identified to species level, whenever possible. Trees were assigned crown class following the classification of Smith et al. (1997).

**Figure 2:**
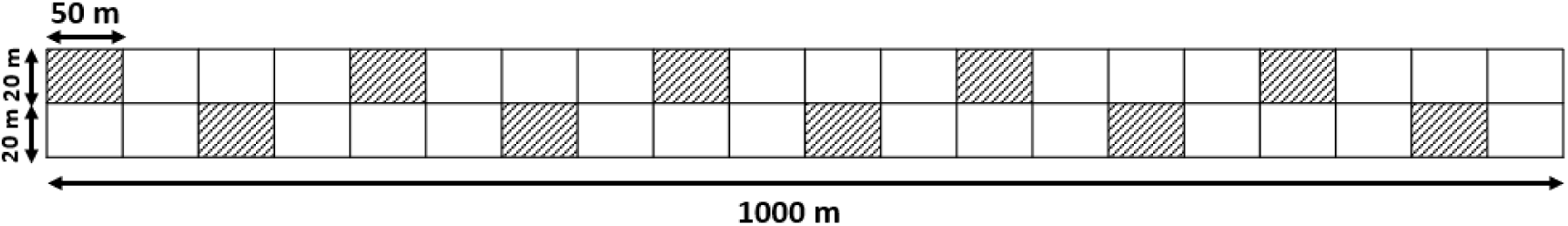
Layout of sampling plot for the study (figure is for representation only and not to scale; filled grids represent the sampled quadrats).

To assess the density and diversity of TMHs and estimate the stand characteristics, all the trees in the selected quadrats were identified and searched for the occurrence of TMHs following Gradstein et al. (2003) and Wolf et al. (2009). All the trees were thoroughly observed from base to top with the help of binoculars (Bushnell legend ultra hd 8×42) by moving around the tree while making observations (Winter and Moller, 2008; Michel and Winter, 2009). The observations were restricted mostly to 10 m height due to lack of visibility or clarity for some of the TMHs such as epiphytes and cavities. However, whenever visible, TMHs at higher locations were also recorded. The trees were vertically divided into 6 zones (Figure 3). Occurrences of all the TMHs were recorded along with the information of the location on the tree – butt, trunk, and branch (primary, secondary, tertiary). Tree species were identified based on standard floras (Hooker, 1897; Manilal, 1988; Vajravelu, 1990; Gamble, 1997).

**Figure 3:**
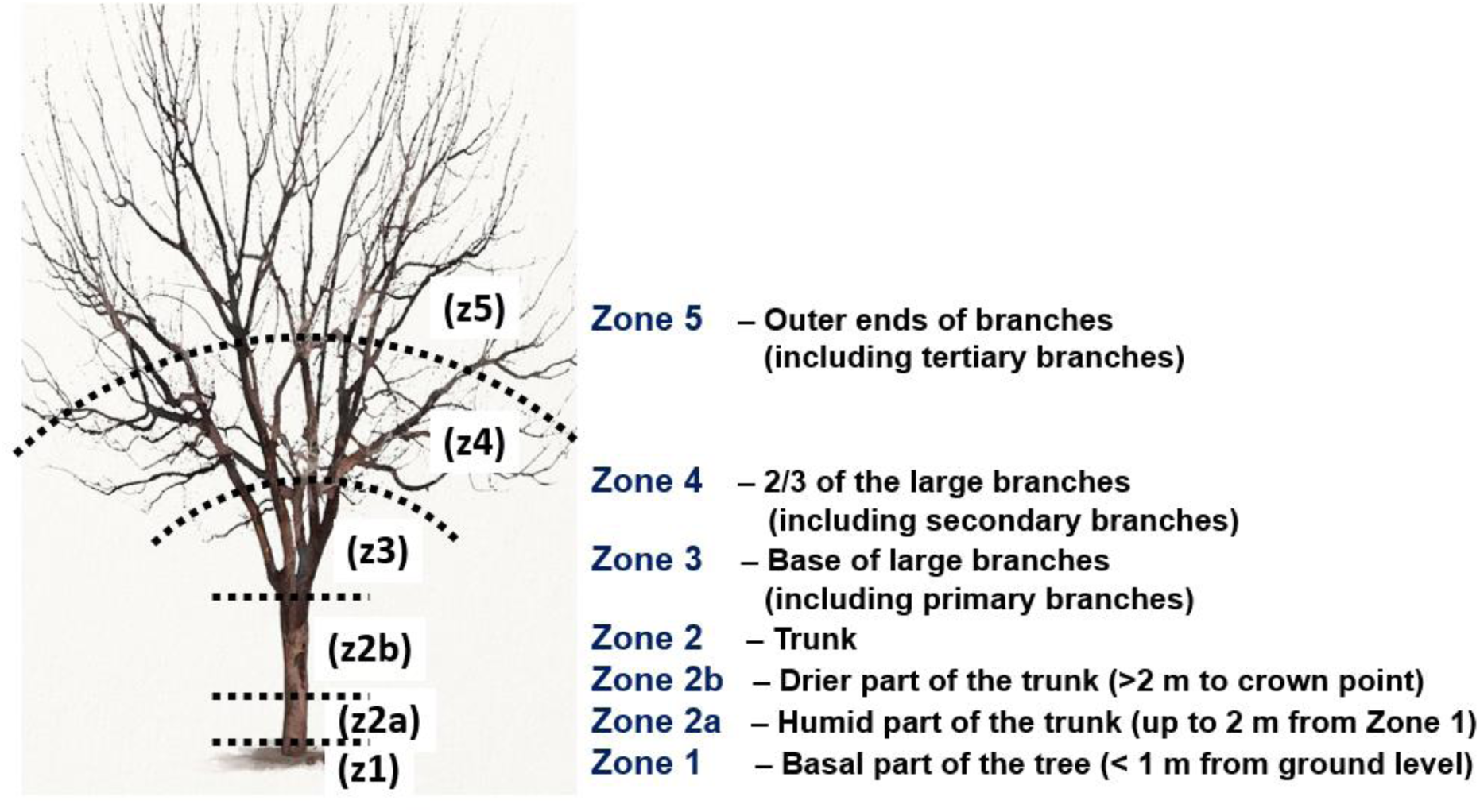
Layout of sampling vertical distribution of tree microhabitats (modified from Gradstein et al., 2003).

### 2.3. Identification and typification of tree microhabitats

The definition of ‘tree-microhabitats’ adopted by Winter and Möller (2008), Vuidot et al. (2011), Larrieu and Cabanettes (2012), Larrieu et al. (2012, 2014, 2018) was followed. For typification of tree microhabitats, we modified the classification and definitions given by Kraus et al. (2016) and Larrieu et al. (2018), and developed a catalogue of tree microhabitats in the study sites. All the microhabitats linked to living trees and snags in the study plots were recorded along with their characteristics.

### 2.4. Characteristics of tree microhabitat host trees

Species and characteristics of the TMH host trees were determined to identify the major host and conditions leading to the occurrence of a single or a combination of TMHs. For each TMH host tree, following characteristics were recorded – the name of the tree species; family; height (m); diameter at breast height (dbh) (m); vitality (live, dead); healthy or unhealthy (in case of live trees). In case of unhealthy trees, cause of unhealthiness was recorded in three broad subcategories – termite (visible termite galleries), fungal decay (fungal fruiting bodies visible on tree) and other (visible symptoms of any other kind of damage/decay). Based on the observations in the field, 76 categories of other causes of unhealthiness were listed (Appendix A). When it was not possible to diagnose and assign a suitable cause of unhealthiness it was listed as unknown. Dead trees were recorded as standing (snag) or fallen/log (downed wood) and analysed separately from the other TMHs present on the standing trees. Tree dimensions were measured for both snags and fallen deadwood. For all the TMHs, every single presence was counted as an occurrence and was summed up to count the total occurrence of each TMH on the tree. Abundance of TMHs were calculated by adding up the total occurrences of each TMH type in each vertical zone of the tree. Quantification of the TMHs on a tree was done considering each vertical zone as a unit of observation and the counting of occurrences varied for each category considering the characteristics of the TMH (Table 1). The existing updated list of TMHs (Kraus et al., 2016) is already exhaustive and includes all the possible types and subtypes of tree microhabitats which may occur on any tree. It includes 15 groups and 47 subgroups of TMHs. Therefore, no further addition to the categories was made. Sub-categories related to other primary excavators (excavated cavities by barbets with cavity entrance > 5 cm, string cavities by barbets) and tree growth form (flute) were added based on field observations along with their characteristics.

**Table 1:**
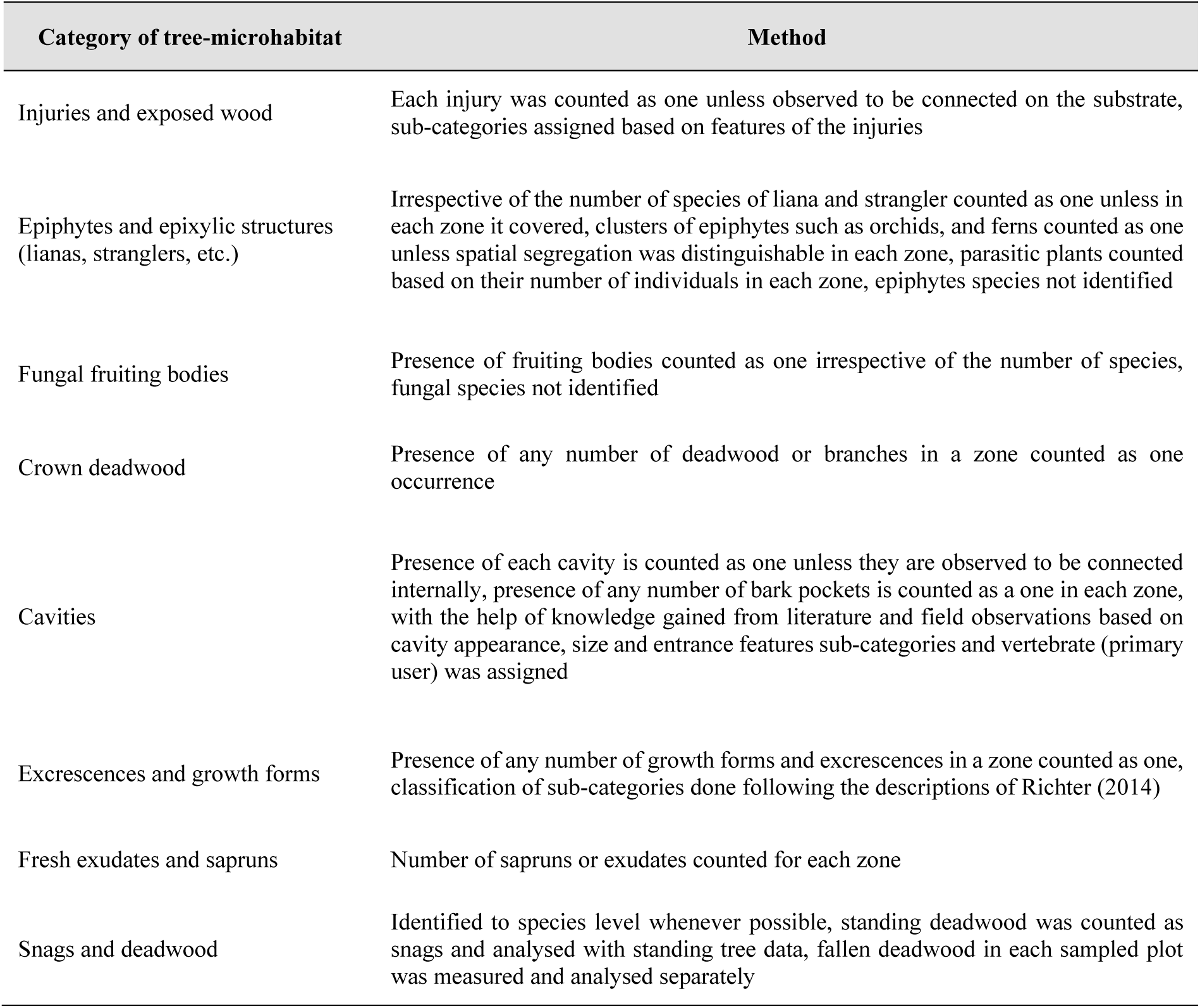
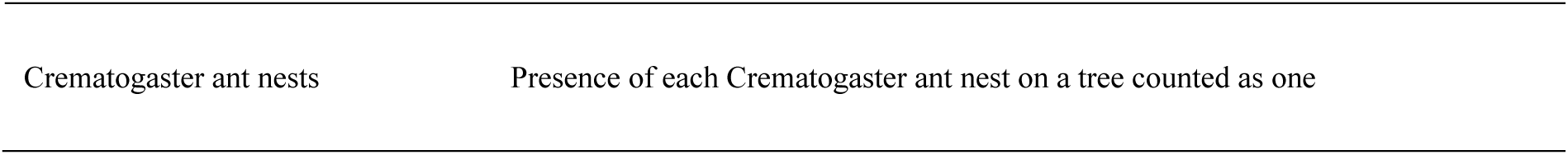
Methods of quantification of the tree-microhabitats on trees.

### 2.5. Analysis

Stand characteristics of the studied habitat types were estimated at the transect level (per hectare, here onwards per ha). The stand-level estimations such as diameter (D), height (H) and basal area (G) (equations in Appendix B) were measured or calculated for all the trees. Density was estimated for all the trees, TMH host trees, cavity host trees, snags, and TMHs (overall and category-wise). All the estimations were carried out for individual trees.

Diversity indices were estimated using standard formulae for all the recorded trees, TMH host trees, cavity host trees and snags. For TMHs (overall and category-wise) frequency (%), density (counts per hectare) and abundance were calculated at the units - per tree and per hectare (Curtis and Mcintosh, 1950). The diversity estimates were calculated following Magurran (2003). Microhabitat Diversity Index was calculated for nine TMH categories as well as 33 sub-categories following Paillet et al. (2018) (equations in Appendix C).

Basic descriptive analysis of numerical data, summaries at plot and transect level, basal area, density, frequency, abundance and microhabitat diversity index calculations were done using basic functions and the Data Analysis Tool Pack of MS Excel. The diversity estimates for host trees except microhabitat diversity index were done using PAST software (Hammer et al., 2001). The stand-level estimates of horizontal and vertical forest structure and composition were done using forest_structure function of ‘forestmangr’ package (Braga et al., 2021). The categorical variables were tested using the Chi-square test with the R package ‘MASS’ (Ripley, 2013). Since the dataset is very large it was assumed that the variable diameter at breast height (dbh), basal area (G) and height (ht) follow normal distribution and were analysed using parametric and non-parametric tests for hypothesis testing. Comparison of mean of numerical variables, occurrence of TMH host trees, occurrence of snags, occurrence of TMHs and significance of variation in occurrences due to stand and tree characteristics in the studied habitats were done using *t* test, ANOVA, Mann-Whitney U test and Kruskal Wallis test in R Software. In case of significant, differences to compare all the possible combinations for levels post-hoc analysis was done using the Dunn Test with the Benjamin-Hochberg method which automatically corrects the ties. Dunn test does pair-wise comparison of all possible pairs following the Holm method for *p*-value adjustment (Dinno, 2015). A comparison of the proportions of categorical variables was done using the Chi-squared test. Graphs and charts for visualization of data were prepared using MS Excel Bluesky Statistics Software V.7.5 (BlueSky Statistics LLC, Chicago, ISA) and R Statistical Software (Version 4.4) (R Core Team 2021). The categorical data is presented in proportions and numeric data as mean±SD, results were considered as significant when *p* < 0.05. Throughout the paper, the numbers in parentheses refer to the number of observations/occurrences unless specifically mentioned.

## 3. Results

### 3.1. Diversity of tree microhabitats

The tree species and tree microhabitat host trees were similar in both the evergreen sites sampled (Figure 4) and hence they are pooled for further analysis.

**Figure 4:**
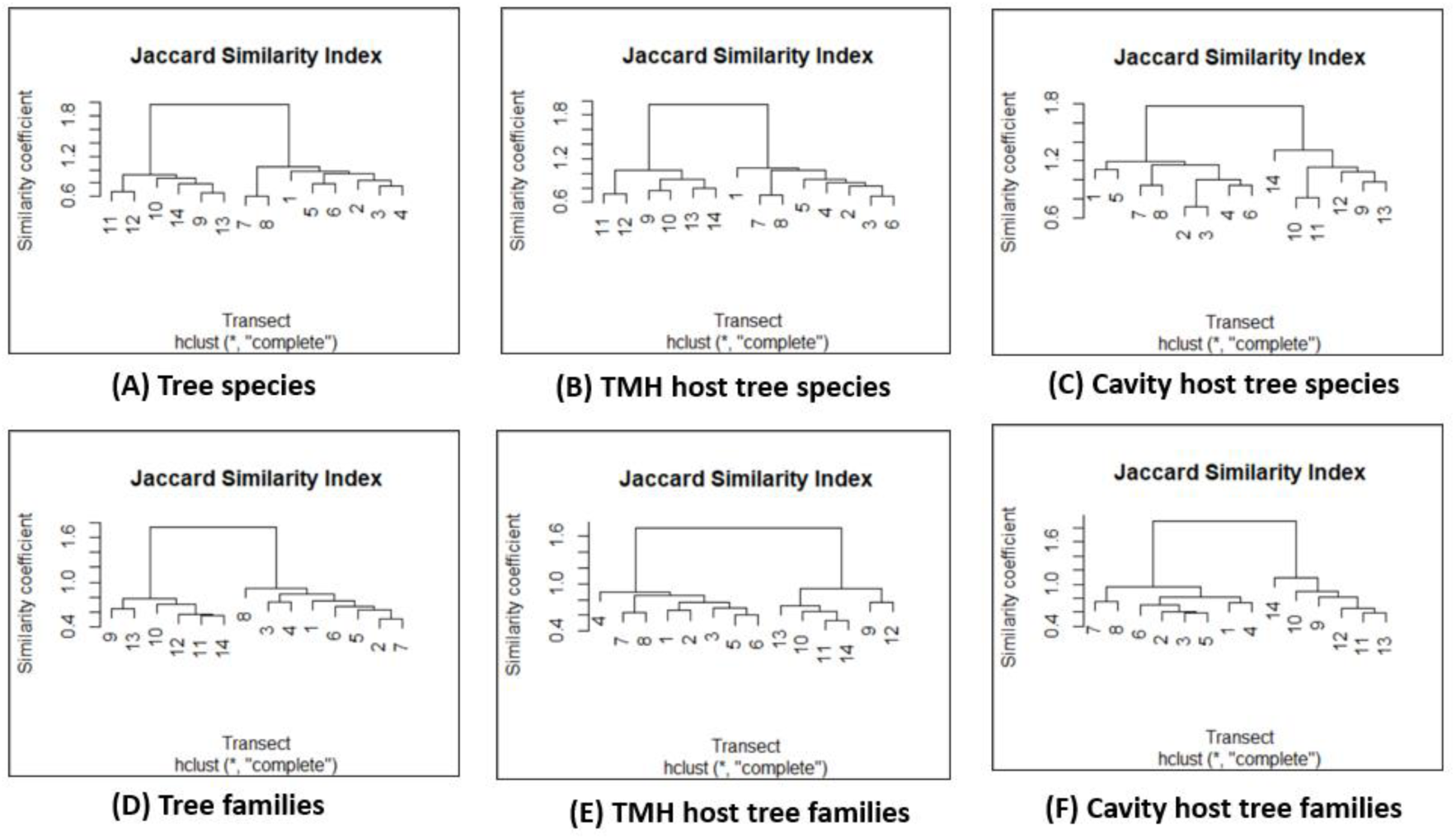
Similarities in the tree species composition of studied transects (clusters indicate similarities in transects: transect 1 to 6 in evergreen forests of SVNP, transects 7 and 8 in evergreen forests of MRF, transects 9 to 14 in PVWLS).

The types of microhabitats were identified and were grouped into 33 sub-categories under nine categories (Table 2). Among these, injuries and exposed wood (m1), excrescences and growth forms (m6) and epiphytic and epixylic structures (m2) were the most diverse TMH categories with nine, eight and seven sub-categories, respectively.

**Table 2:** Typification of tree microhabitats, definition and characteristics recorded.

| TMH category* | Definition | TMH sub-category* | Definition | Characteristics |
| --- | --- | --- | --- | --- |
| m1 | Injuries and Exposed Woods | m1_bt | Broken tree top | Presence, Occurrences (Abundance) |
|  |  | m1_bb | Broken fork and branches |  |
|  |  | m1_fi | Fissures |  |
|  |  | m1_cr | Cracks and splits |  |
|  |  | m1_ss | Splintered stem |  |
|  |  | m1_an | Animal scratch/peels |  |
|  |  | m1_fd | Fire damage |  |
|  |  | m1_te | Termite decay |  |
|  |  | m1_un | Unknown source of rot or decay |  |
| m2 | Epiphytic and Epixylic Structures | m2_sf | Strangler Ficus | Presence, Occurrences (Abundance) |
|  |  | m2_li | Climbers and lianas |  |
|  |  | m2_lc | Lichens |  |
|  |  | m2_br | Bryophytes |  |
|  |  | m2_pt | Pteridophytes |  |
|  |  | m2_or | Orchids |  |
|  |  | m2_pe | Other epiphytes |  |
| m3 | Fungal Fruiting Bodies | m3_ffb | Fruiting bodies of rot fungi | Presence, Occurrences (Abundance) |
| m4 | Crown Deadwood | m4 | Dead branches on the stem and in the canopy | Presence, Occurrences (Abundance) |
| m5 | Cavities | m5_cv | Number of cavities (natural and excavated) | Presence, Occurrences (Abundance), Dimensions, Orientation |
|  |  | m5_na | Number of natural cavities |  |
|  |  | m5_ex | Number of excavated cavities |  |
|  |  | m5_bp | Bark-pockets |  |
| m6 | Excrescences and Growth Forms | m6_bu | Buttress | Presence, Occurrences (Abundance) |
|  |  | m6_fl | Flutes |  |
|  |  | m6_se | Seams |  |
|  |  | m6_cs | Crooked stem |  |
|  |  | m6_fo | Forked stem/forking |  |
|  |  | m6_sp | Spiral/twisted stem |  |
|  |  | m6_ca | Cankers |  |
|  |  | m6_bl | Burls |  |
| m7 | Deadwood – snags, stumps, fallen deadwood | m7_st | Snags | Presence, Occurrences (Abundance), Dimensions |
|  |  | m7_fl | Fallen deadwood |  |
| m8 | Fresh Exudates | m8 | Sapruns | Presence, Occurrences (Abundance) |
| m9 | Crematogaster Ant-nests | m9_ca | Crematogaster ant-nests | Presence, Occurrences (Abundance) |

\* The category and sub-category names indicate the codes used for analysis and in text for referring to respective TMH.

The average diversity of the nine types of TMHs (averaged from per transect) in evergreen was 79.31 %, and in deciduous it was 74.44 %, but not differ significantly (One-way ANOVA *F* = 1.374, *p* = 0.2638). The most abundant TMH type in the evergreen forests was excrescences and growth forms (n = 1,652), followed by epiphytic and epixylic structures (n = 1,638) and then by injuries and exposed wood (n = 1,331). In deciduous forests, the most abundant TMH was injuries and exposed wood (n = 633), followed by epiphytic and epixylic structures (n = 396), and then by crown deadwood (n = 392). The average diversity of the 33 TMH sub-categories (averaged per transect) was 54.47 % in evergreen and 52.37 % in deciduous. When combined, the difference in the diversity of the TMH subcategories was not significant (*F* = 0.0374, *p* = 0.8499). Further, among the sub-categories in the evergreen and deciduous stands, the difference in abundance was significant only for cankers (Levens test for equality of variance: *F* = 12.807, *p* < 0.01) and standing deadwood (*F* = 10.227, *p* < 0.01). Among the 33 sub-categories, in the evergreen forests, the most abundant were burls (n = 1,482), followed by bryophytes (n = 743), and then by lianas (n = 654). In the deciduous forests, the most abundant TMH sub-categories were crown deadwood (n = 403), followed by cavities (n = 304), and then by lianas (n = 300) (Figure 5).

**Figure 5:**
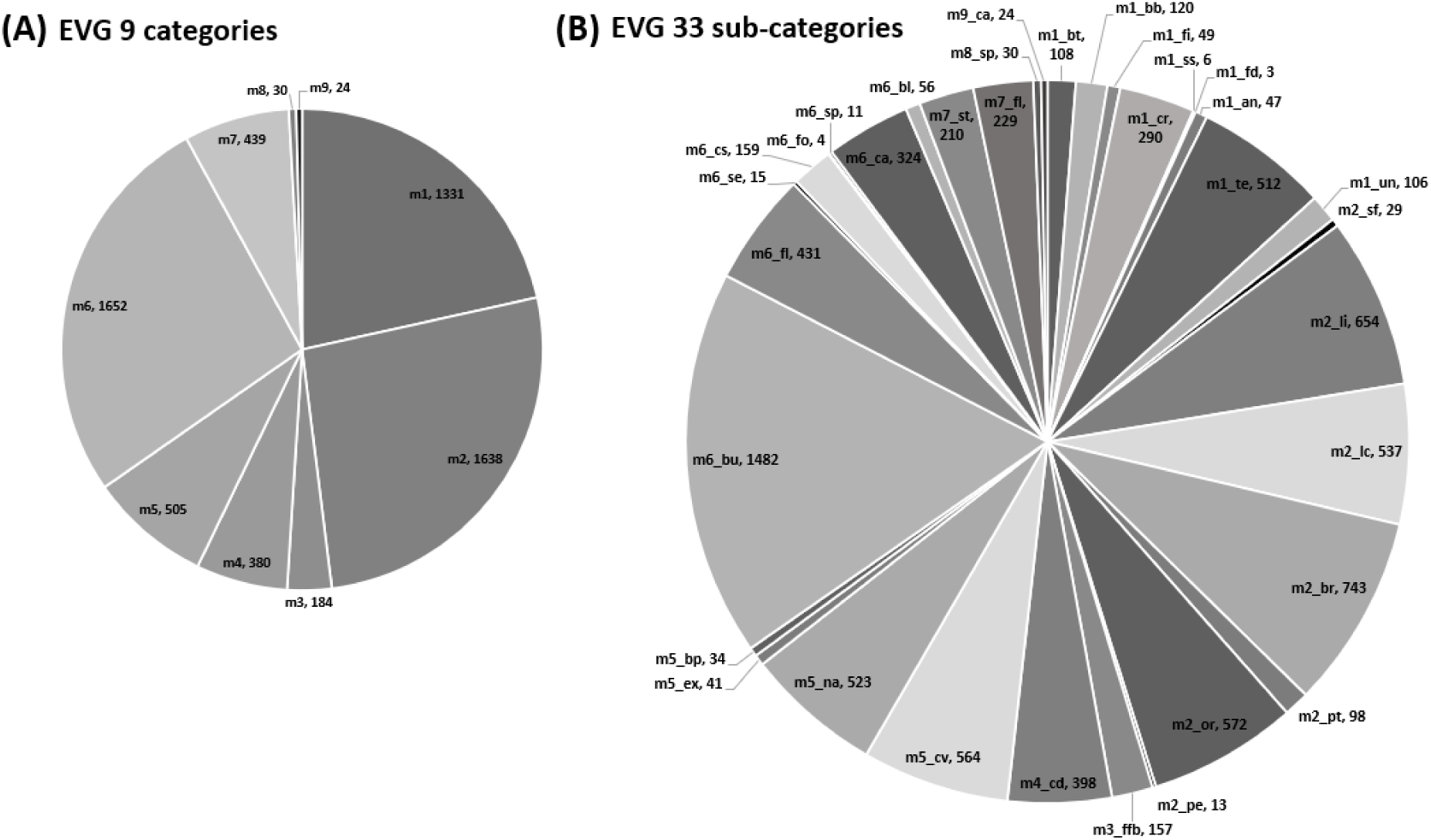
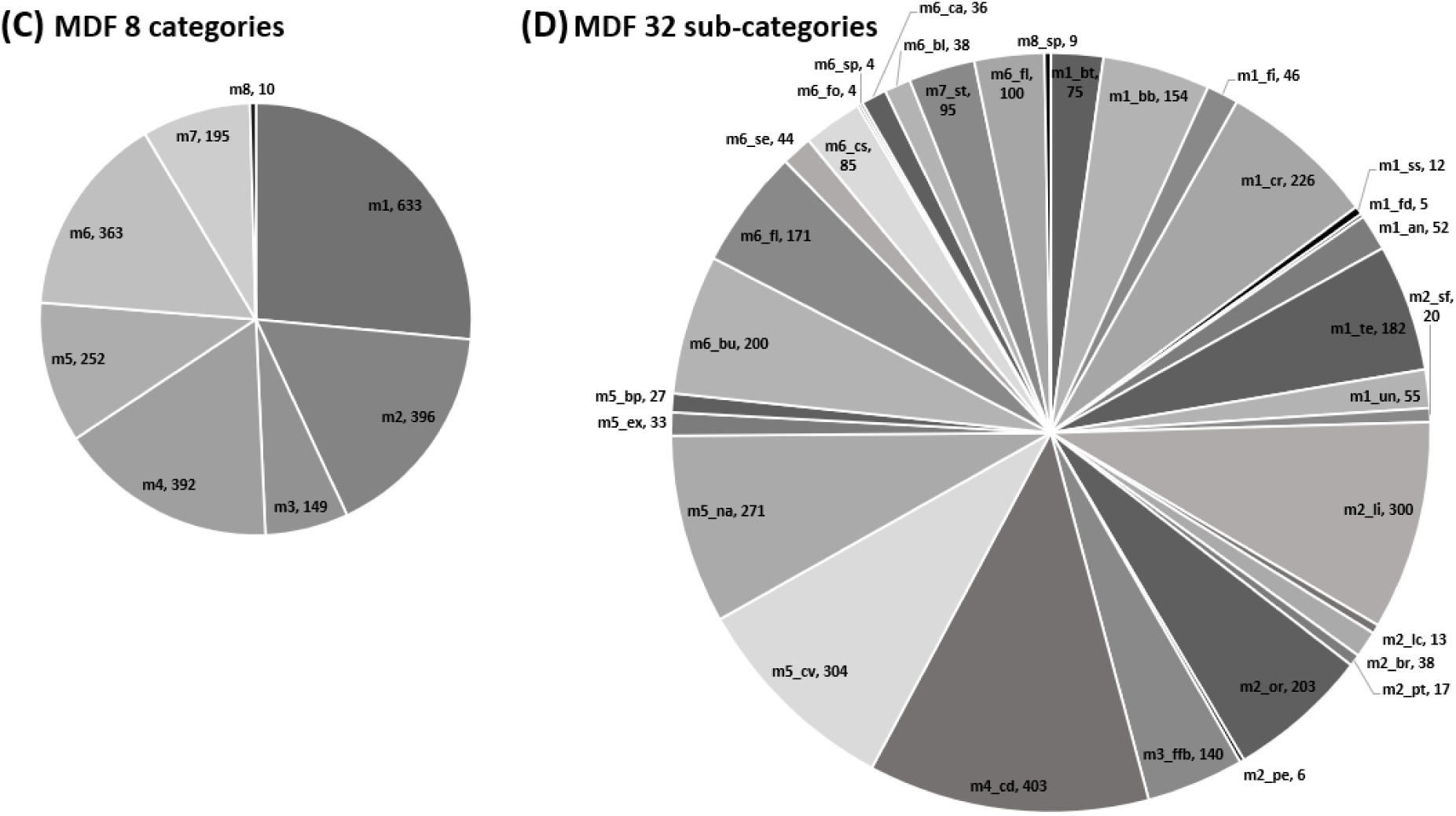
Proportion of TMH types occurring in the studied habitats: (A) 9 types in evergreen habitat; (B) 33 sub-categories in evergreen habitat; (C) 8 types in moist deciduous habitat; (D) 32 sub-categories in moist deciduous habitat.

### 3.2. Horizontal forest structure and distribution of tree microhabitats

#### 3.2.1. Structure and composition of host species

In evergreen habitats, a total of 6,363 trees were sampled which belong to 168 species (including 37 unidentified), 116 genera (2 unidentified) and 51 families. The trees in evergreen habitats had a mean dbh of 0.26±0.20 m (range: 0.10 to 2.31 m) and a mean height of 13.90±6.44 m (range: 1 to 40 m). The density of overall trees was 795.4±110.6 ha^-1^, the density of trees hosting at least one TMH was 454.6±115.1 ha^-1^, the density of snags was 54.9±19.7 ha^-1^ and trees hosting at least one cavity was 56.3±11.3 ha^-1^. In evergreen, 3,637 trees were observed to host 7,782 TMHs with a density of 12.4±4.9 TMHs per tree and 972.8±341.4 TMHs ha^-1^ (Table 3). The TMH host trees in this habitat belong to 105 species (34 unidentified) in 100 genera (2 unidentified), and 50 families. The key tree species hosting microhabitats were *Palaquium ellipticum* (368 individuals), *Cullenia exarillata* (313), *Dimocarpus longan* (292), *Drypetes venusta* (174), *Cryptocarya wightiana* (154) and representative families were Lauraceae (416), Malvaceae (339), Sapotaceae (375), Myristicaceae (332), Sapindaceae (296). The cavity host trees belong to 74 species (22 unidentified), 56 genera (1 unidentified), and 35 families. The major cavity host species were *Cullinea exarillata* (50), *Myristica fatua* (46), *Drypetes venusta* (42), *Mesua ferrea* (30), *Dimocarpus longan* (29), *Elaeocarpus tuberculatus* (27). The major families hosting cavities were Myristicaceae (72), Malvaceae (52), Lauraceae (46), Putranjivaceae (43), Calophyllaceae (35). Of the 6,363 trees sampled in evergreen habitats, 134 live trees could not be identified up to species level and 145 dead individuals could not be identified as they were in advanced decay stages. These included 87 live TMH host trees belonging to 45 taxa and 143 snags, and seven live cavity host trees belonging to seven taxa.

**Table 3:**
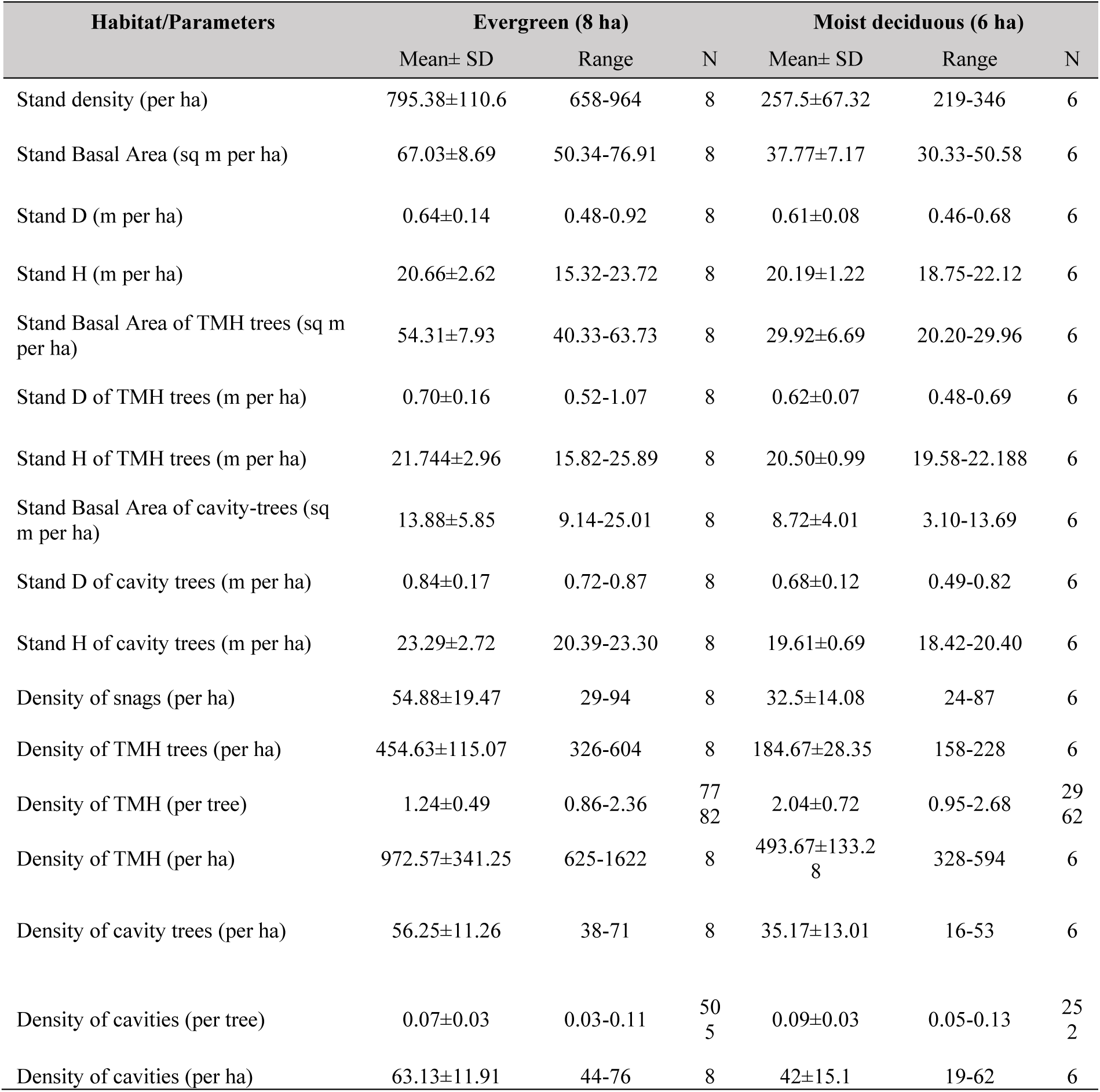
Summary of the composition in the studied habitats.

In deciduous habitat, 1,545 trees were sampled belonging to 74 species (5 unidentified), 56 genera (2 unidentified) and 39 families. The taxa of two live individuals and 13 dead individuals out of 1,545 could not be identified. The mean dbh of trees was 0.35±0.21 m (range: 0.10 to 2.31 m) and the mean height was 15.05±7.01 m (range: 0.70 to 35.00 m). In deciduous habitat, the density of trees was 257.5±67.3 ha^-1^, the density of trees hosting at least one TMH was 184.7±28.4 ha^-1^, the density of snags was 32.5 14.1ha^-1^, and the density of trees hosting at least one cavity was 35.2±13.0 ha^-1^. In this habitat, 1,108 trees were observed to host 2,962 TMHs with a density of 20.4±7.2 TMHs per tree and 493.7±133.3 TMHs ha^-1^. The TMH host trees belong to 70 species (2 species unidentified), 50 genera (2 unidentified), and 35 families. The representative TMH host trees species were *Xylia xylocarpa* (282 individuals), *Terminalia paniculata* (105), *Dillenia pentagyna* (82), *Tectona grandis* (61), *Macaranga peltata* (57) and families Leguminosae (325), Combretaceae (161), Euphorbiaceae (93), Malvaceae (86), Dilleniaceae (83). The cavity host trees belong to 44 species (2 unidentified), 39 genera, and 27 families. Four snags could not be identified to the species level. The major cavity host species were *Xylia xylocarpa* (67), *Terminalia paniculata* (35), *Dillenia pentagyna* (27), *Cleistanthus collinus* (17), *Macaranga peltata* (10). The major families hosting cavities were Leguminosae (74), Combretaceae (38), Dilleniaceae (27), Euphorbiaceae (22), and Malvaceae (14).

Across the habitats, the density of TMH host trees was very high compared to the cavity host trees (Table 3). The representative host tree species in both habitats were also the species with high relative frequency, relative density and relative dominance and with high area coverage (Table 4). Proportionate to the stand composition, the proportion of TMH and cavity host trees was high in the deciduous habitat (71.72 % and 13.72 %, respectively) compared to the evergreen habitat (57.16 % and 7.36 %, respectively). However, the number of TMHs was high in the evergreen habitat (n = 7,782) compared to deciduous (n = 2,962).

**Table 4:**
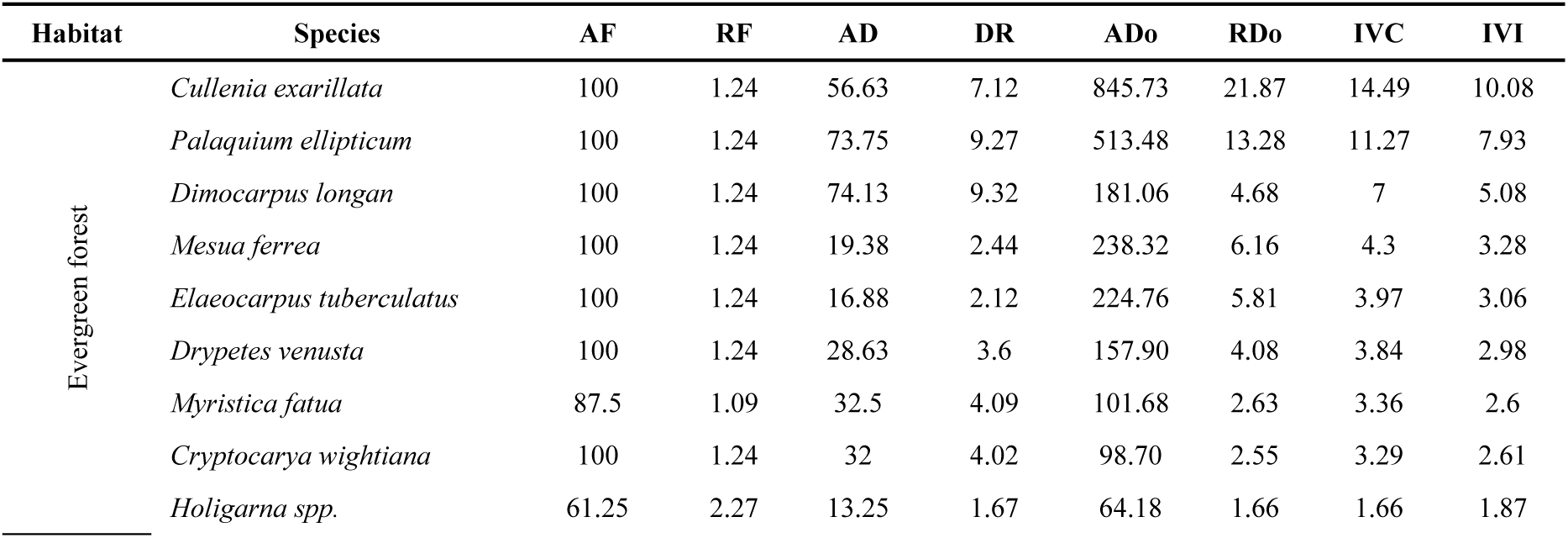
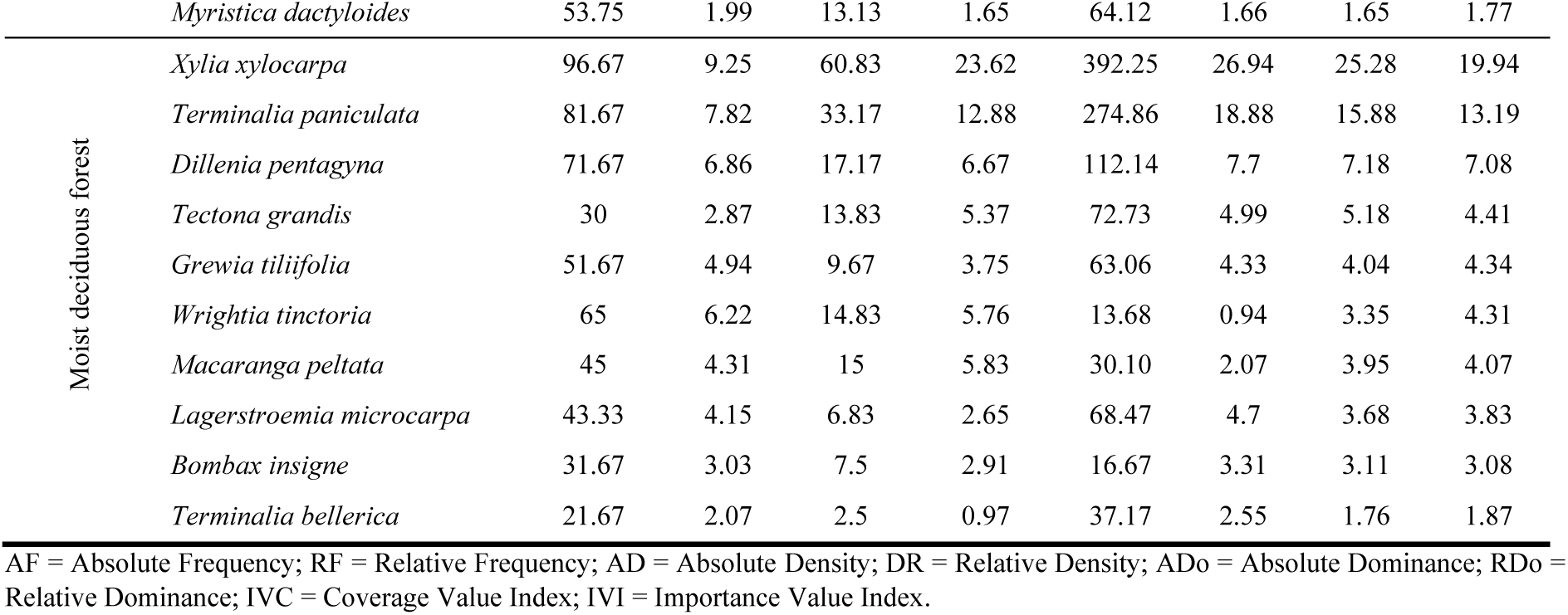
Density estimates of the top 10 tree microhabitat-bearing tree species in evergreen and deciduous habitats.

In evergreen habitat, majority of the trees and TMH host trees were in the height class 6-16 m and dbh class 0.1-0.15 m (Figure 6 A and B). Majority of the trees were in large diameter classes, and the average basal area in the habitat was 67.03±37.77 sq m ha^-1^. A major proportion of the trees and TMH host trees were live healthy trees (73.44 % and 55.82 %), but the major proportion of the cavity host trees were live unhealthy trees (72.28 %). There was a significant difference between the dbh of TMH host trees in live healthy, live unhealthy and snag vitality classes in the transects (ANOVA, *F* = 23.638; *p*<0.001) (Table 5). The post-hoc analysis showed that the dbh of snags and healthy TMH host trees were significantly different (*p* < 0.05). The difference in the diameter of snag and unhealthy trees (*p*<0.001) and healthy and unhealthy trees (*p*<0.001) were highly significant. Among the cavity hosts, the difference in the dbh of live healthy, live unhealthy trees and snags were not significant (*p* = 0.4623). The difference in the height of TMH host trees was also highly significant (*F* = 132.042; *p*<0.001). Post-hoc analysis shows that height of unhealthy and healthy trees was significant (*p* < 0.05), and snags and healthy trees (*p*<0.001) and unhealthy trees was highly significant (*p*<0.001). The difference in the height of the cavity hosts were estimated to be highly significant (*F* = 15.281; *p*<0.001) except for the height of healthy and unhealthy trees (*p* = 0.5723). The difference in the basal area of TMH hosts were also highly significant (*F* = 23.406; *p*<0.001), but post hoc analysis shows that the difference in the basal area of healthy trees and snags was not significant (*p* = 0.9339). The difference in the basal area of the healthy and unhealthy trees (*p*<0.001) and unhealthy trees and snags (*p* < 0.01) hosting TMHs was highly significant, but difference in the basal area of trees hosting cavities was not significant (*p* = 0.6014). In evergreen habitats, the frequency of TMH host trees ranged from 43 % to 88.05 % and cavity host trees ranged from 5.23 % to 8.97 % in different transects. Health of the tree is observed to influence the abundance of TMHs and cavities. Unhealthy trees and snags had high occurrences of TMHs and healthy trees had more cavities.

**Figure 6:**
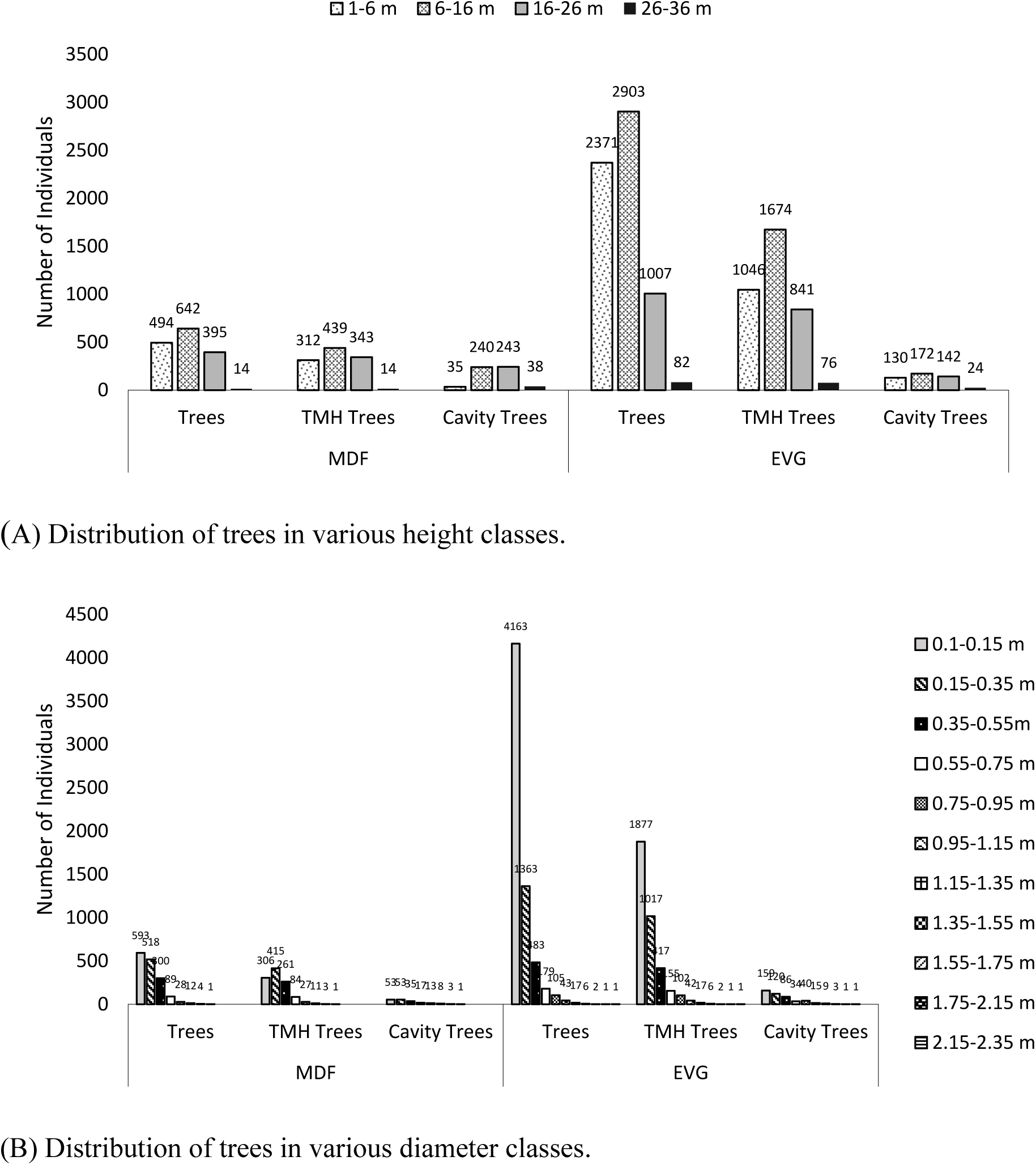
Distribution of trees in various height and diameter classes in the studied habitats.

**Table 5:**
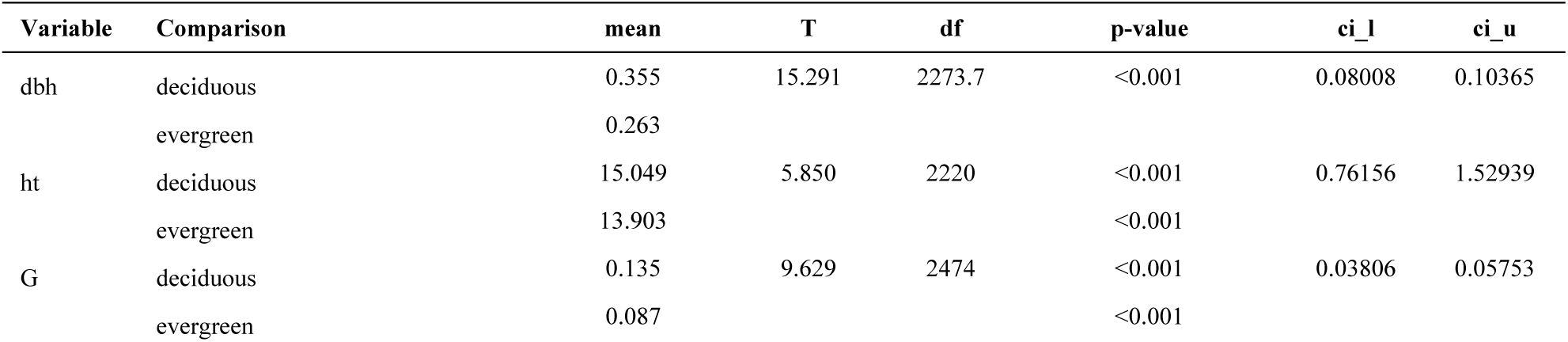
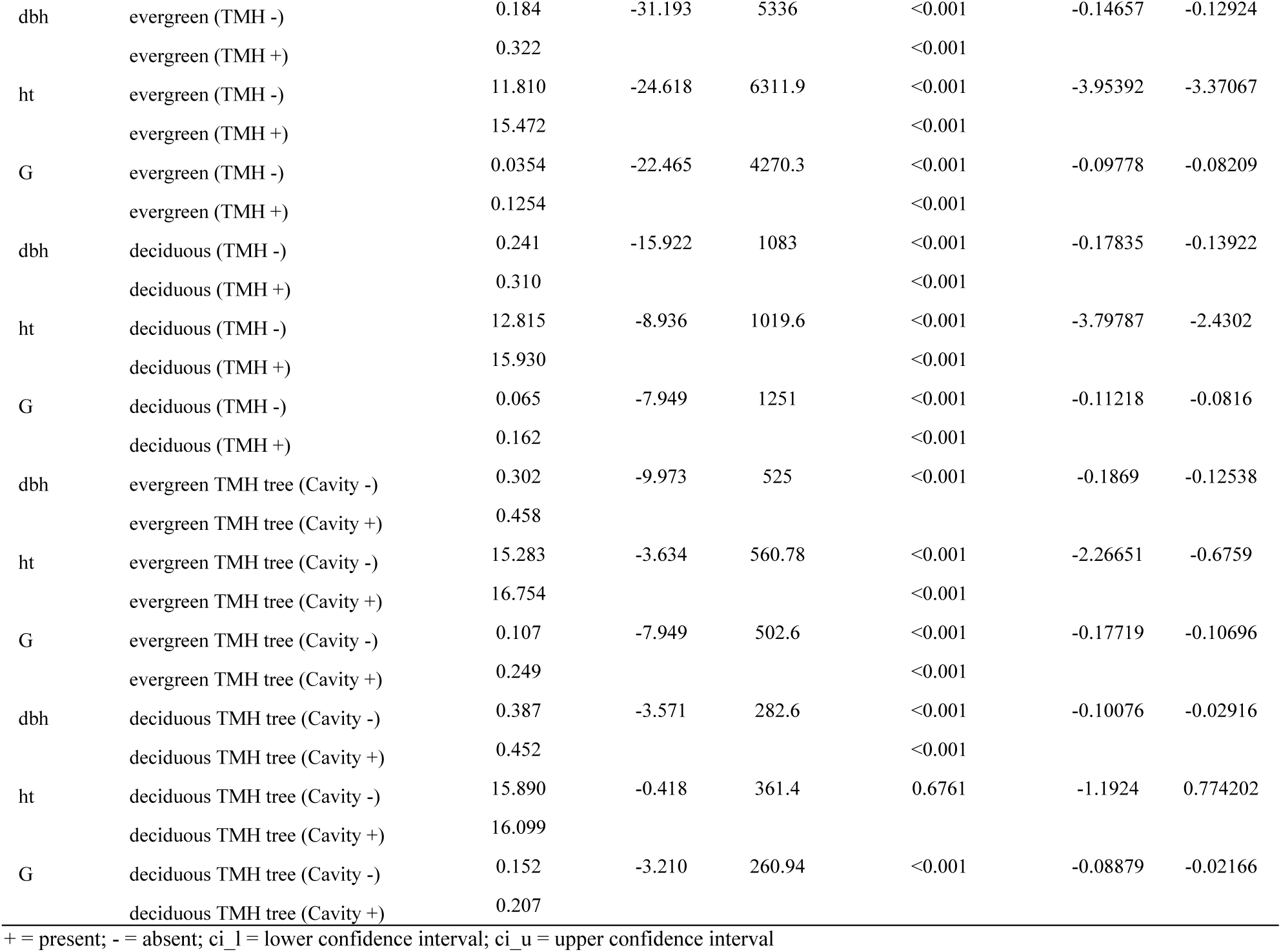
Comparison of means of stand characteristics of evergreen and moist deciduous habitat – diameter at breast height (dbh), height (ht) and basal area (G).

In deciduous habitat, among the sampled individuals, most of the trees were healthy (54.24 %), but microhabitats and cavities were more on unhealthy host trees (54.33 % and 73.02 % respectively). The majority of the trees and TMH host trees were in the height class 6-16 m. In general, trees were in the dbh class 0.1-0.15 m, but the majority of TMH and cavity host trees were in the dbh class 0.15-0.35 m (Figure 6 A and B). The average basal area estimated in the habitat was 37.77±7.17 sq m ha^-1^. The diameter of TMH host trees was significantly different among the healthy, unhealthy trees and snags (*F* = 7.352; *p*<0.001), but the difference in the dbh of healthy and unhealthy trees was not significant (*F* = 0.017; *p* = 0.2163). The dbh of cavity host trees in the three vitality classes did not differ significantly (*F* = 0.004; *p* = 0.9956). The difference in the height of the TMH host trees in different vitality classes was highly significant (*F* = 63.923; *p*<0.001). Post-hoc analysis also shows that the differences in height of the TMH host trees was significant for the live healthy and unhealthy (*p*<0.001), live healthy tree and snags (*p*<0.001), and unhealthy and snags (*p*<0.001). The height of the cavity host trees in the three vitality classes differed significantly (*F* = 6.409; *p* < 0.001) respectively. However, the difference in the height of healthy and unhealthy cavity host trees was not significant (*p* = 0.2133). There was highly significant difference in the height of cavity host trees in the vitality class snags and healthy tree (*p* < 0.001), and snags and unhealthy trees (*p* < 0.01). The differences in basal area of TMH host trees in different vitality classes were significantly different (*F* = 3.919; *p* < 0.05), but the post hoc analysis revealed that the difference between the healthy and unhealthy trees were not significant (*p* = 0.4187), but there was significant difference between basal area of snags and healthy trees (*p* < 0.05), and snags and unhealthy trees (*p* < 0.05). The differences in the basal area of cavity host trees which were in the vitality classes healthy, unhealthy and snag were not significant (*p* = 0.8955). The healthy trees tend to have higher counts of cavities, but unhealthy trees and snags tend to have more occurrences of TMHs.

The Chi-squared test shows that the vitality of trees, cause of unhealthiness of trees, decay class of snags, and presence and absence of microhabitats including cavities differed significantly between the evergreen and deciduous habitats (Table 6). Even within the habitats, the stand heterogeneity in the transects resulted in significant variation in TMH occurrences (evergreen: *χ^2^* = 1353.9, *p* <0.001; deciduous: *χ^2^* = 464.07, *p* <0.001; Table 6).

**Table 6:** Results of chi-squared test for comparing the effect of habitat on the occurrence of trees in different vitality classes, host individuals for tree microhabitats and cavities, and presence of tree microhabitat and cavity.

| Variable 1 | Variable 2 | Chi-squared ($\chi^2$) | df | p-value |
| --- | --- | --- | --- | --- |
| Habitat | Vitality | 217.57 | 2 | <0.001 |
| Habitat | Cause of Unhealthy Tree | 290.24 | 4 | <0.001 |
| Habitat | Decay class of snags | 23.884 | 5 | <0.001 |
| Habitat | Cavity presence-absence | 63.307 | 1 | <0.001 |
| Habitat | Cavity origin | 69.617 | 3 | <0.001 |
| Habitat | TMH presence-absence | 109.15 | 1 | <0.001 |
| Evergreen | TMH presence-absence | 1353.9 | 2 | <0.001 |
| Deciduous | TMH presence-absence | 464.07 | 2 | <0.001 |

#### 3.2.2. Diversity of tree microhabitat host species

In the evergreen stands, the average species richness was 80±13.77 species, with an average dominance of 0.05±0.01. The Simpson index was estimated to be 0.95±0.01, the Shannon Index was 3.57±0.11, average evenness was 0.45±0.04, equitability 0.82±0.01, Fisher’s alpha was 22.66±5.97, Berger Parker index was 0.12±0.02. Chao-1 was estimated to be 115.41±39.05, indicating the probability that sites may have more species and are missed in samples due to their distribution pattern (Figure 7). The average richness of TMH host trees was 64.38±13.84 species which is less compared to the overall species richness in the area indicating that all the species do not host TMHs. The dominance was 0.05±0.01, which was similar to the dominance observed when non-host species were also included in the analysis. It also indicates that some species are dominant hosts of TMHs. Similarly, evenness (0.51±0.04) and equitability (0.84±0.02) were also estimated to have similar values. Simpson index was 0.95±0.01, similar to the estimate including other species indicating high diversity in the host species. Shannon index was lesser compared to the overall diversity estimate of 3.48±0.13 as the number of host species and therefore the abundance was less compared to all the recorded species. Fisher’s alpha was 21.28±5.54 (less than the estimate for all the species) as a few of the recorded species never hosted any TMH. The Berger-Parker index indicates the dominance of the most abundant species and it was higher than the estimate for all the species (0.13±0.01) probably because the dominant host species were also the most abundant in the sampled sites. The Chao-1 estimate for TMH host species 87.27±29.17 indicates that the richness of host species could be more than 100 species. The average richness of cavity host was 23.50±2.78 species which could be up to 50 species as estimated with Chao-1 (39.55±9.80). The difference in species richness was highly significant between species recorded in the habitats, hosts of TMHs and cavities using Kruskal Wallis test (*H* = 28.605, *p*<0.001). The number of species individuals recorded in the inventory and hosts of TMHs and cavities also varied significantly (*H* = 29.633, *p*<0.001). The relatively higher dominance 0.06±0.01 indicates host specificity for cavities, but the difference in the dominance of species as host of microhabitats and cavities was not significant (*H* = 2.251, *p* = 0.3246). The difference in Chao-1 index values were also highly significant (*H* = 24.935, *p*<0.001), but only for sampled tree species and cavity host (*p*<0.001) and species of tree microhabitats and cavities (*p*<0.001) probably because the hosts species tend to be both dominant. However, difference in the dominance of TMH and cavity host species in evergreen habitats was significant (*H* = 8.82, *p* < 0.05). As all the TMH host trees did not host cavities, the Simpson index 0.94±0.01, Shannon index 3.01±0.15, evenness 0.87±0.06 and Berger Parker index 0.17±0.04 were estimated to have relatively lower values. Relatively high equitability 0.95±0.03 was estimated probably because the cavity hosts were nearly equally abundant and this difference was highly significant (*H* = 25.739, *p*<0.001). Fisher’s alpha for cavity host species 14.37±2.64 was estimated even lesser than the TMH host indicating that all the TMH hosts do not host cavities. The Simpson index remained high for all the species as well as host species in both the habitats, the difference in this index values of the two evergreen and deciduous habitats estimated using Mann Whitney U test were significant (*U* = 117, *p*<0.001). These differences were significant for Shannon index (*U* = 7.974, *p* <0.05), Fisher’s alpha (*U* = 9.067, *p* < 0.05) and evenness (*U* = 31.738, *p* < 0.01) and not for Simpson index (*U* =2.224, *p* =0.329) and Berger-Parker index (*U* = 2.444, *p* = 0.295). The Berger-Parker index did not differ significantly for the inventoried species and host of cavities in the evergreen habitat (Kruskal Wallis test *H* = 6.488, *p* = 0.936) probably because mainly the dominants are hosting TMHs and cavities in both the habitats.

**Figure 7:**
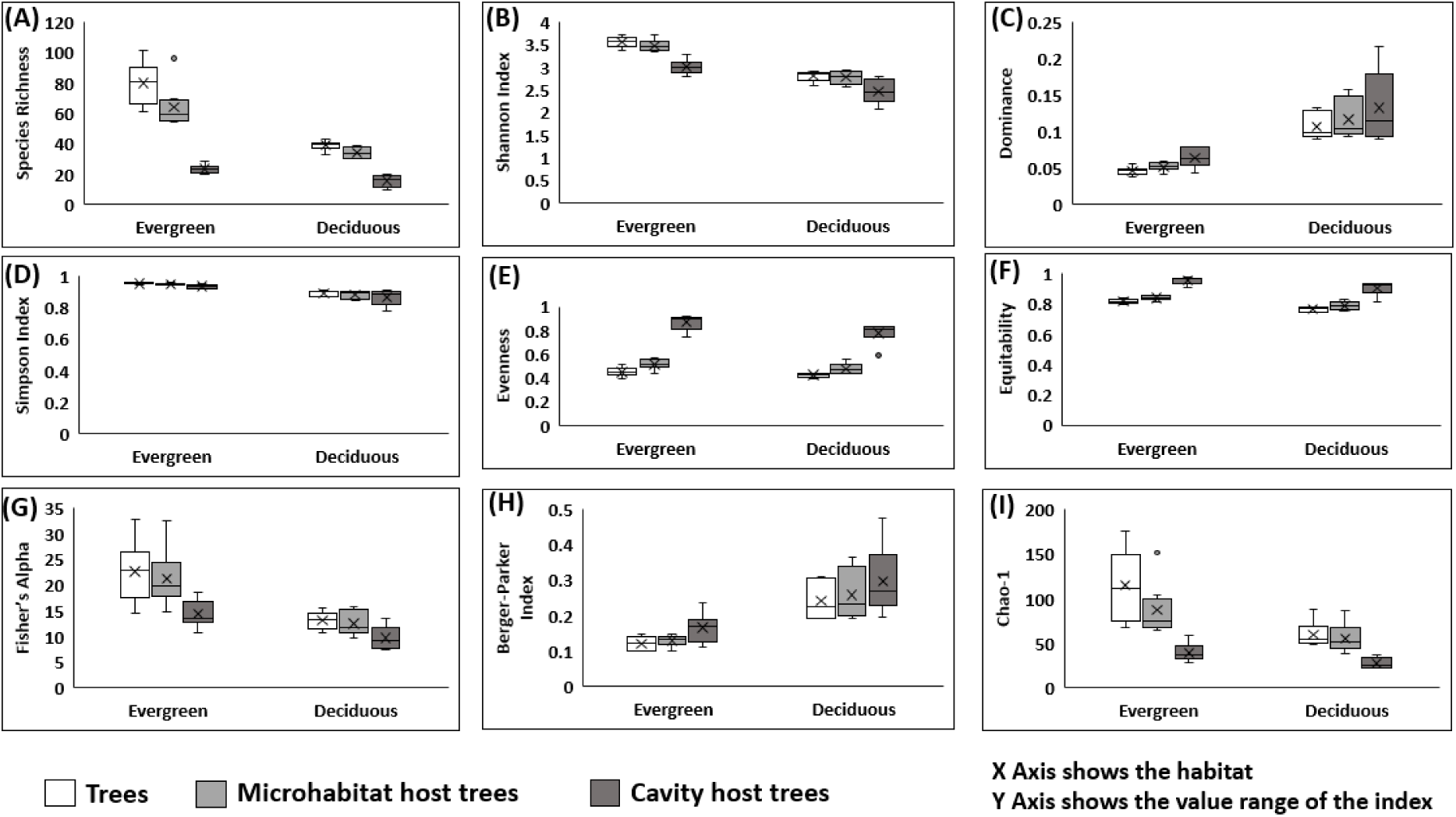
Diversity indices estimated for the studied habitats.

### 3.3. Vertical forest structure and distribution of tree microhabitats

Among the trees sampled in the evergreen habitat, 73.44 % of individuals were healthy, 23.36 % were unhealthy and 3.3 % were dead. 35.01 % of individuals were damaged by termites, 9.10 % were damaged by fungal decay and 55.89 % were damaged by other causes. Among the TMH host trees, 35.04 % were damaged by termites, 9.42 % were decayed by fungi and 55.55 % were injured and/or damaged by other causes. 7.36 % of individuals hosted cavities out of which 50.82 % were damaged by termites, 11.81 % were decayed by fungi and 37.36 % were damaged by other causes. Most of the sampled trees belonged to the intermediate class (40.57 %), followed by codominant (17.90 %), dominant (15.27 %) and emergent (5.73 %). Among the individuals hosting TMH, 49.31 % were intermediate, 22.25 % were overtopped, 15.55 % were codominant, 10.63 % were dominant and 2.26 % were emergent. Among the individuals hosting cavities, 41.01 % were intermediate, 19.08 % codominant, 19.08 % overtopped, 15.13 % dominant and 5.71 % emergent (Figure 8). The differences in the dbh, height and basal area of trees in various crown classes were tested to identify the influence of crown class on TMH presence and abundance. The differences in the dbh of TMH host trees (*F* = 936.981; *p*<0.001) and cavity host trees (*F* = 132.343; *p*<0.001) in various crown classes were highly significant. The differences in the heights of TMH and cavity host trees in various crown classes were highly significant (*F* = 3993.11; *p*<0.001 and (*F* = 618.039; *p*<0.001, respectively). The basal area of TMH and cavity host trees was significantly different among the classes (*F* = 525.537; *p*<0.001) and (*F* = 74.240; *p*<0.001) respectively. However, the post hoc analysis revealed that the difference in basal area of cavity trees in overtopped and intermediate classes was not significant (*F* = 0.075; *p* = 0.0859). The total number of TMHs recorded from the evergreen habitat was 14,846. The highest occurrence of TMHs was recorded in the lower part of the trunk (Zone 2a) (4,028), followed by the base of the tree (Zone 1) (3,011), followed by the upper part of the trunk (Zone 2b) (2,873). In evergreen stands, epiphytic and epixylic structures were most abundant with the highest occurrence in the upper part of the trunk (Zone 2b) (1,305) followed by the primary branches (Zone 3) (1,197), followed by the lower part of the trunk (Zone 2a) (1,096) (Figure 9).

**Figure 8:**
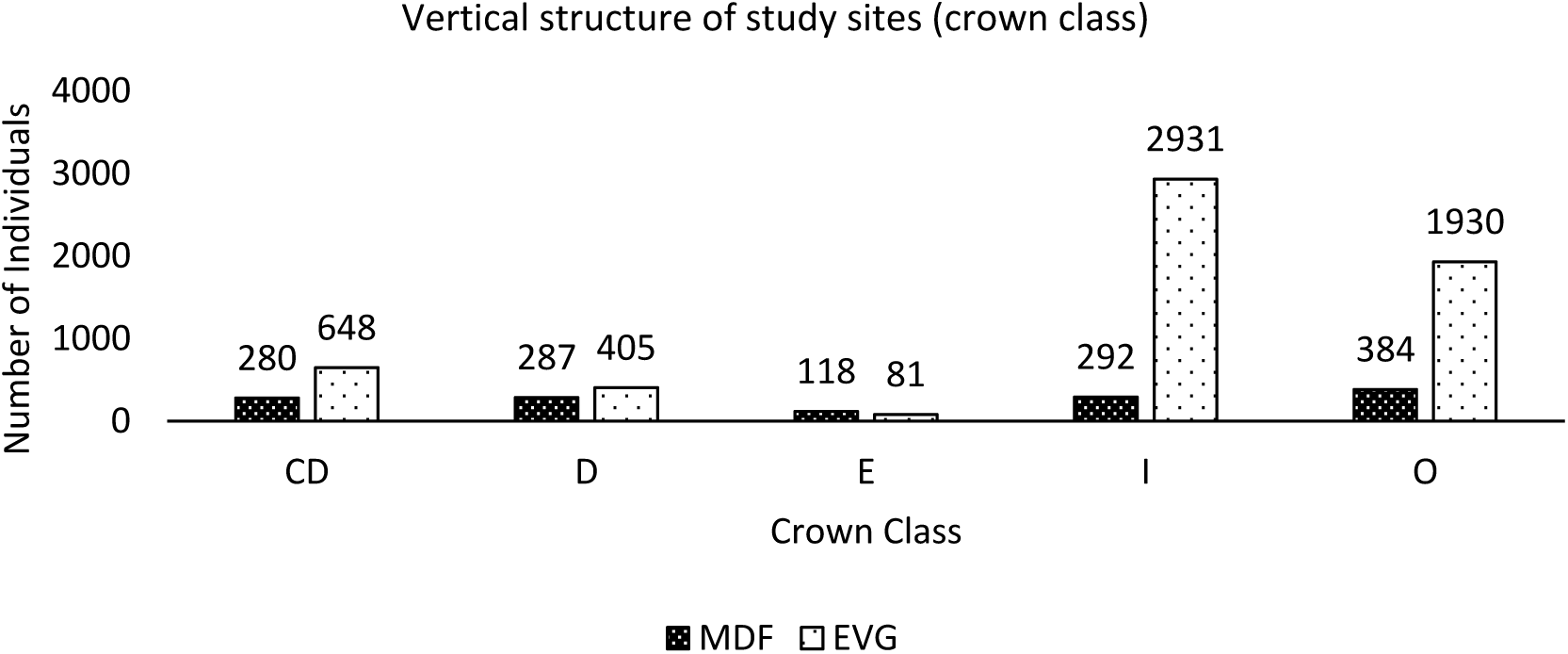
Distribution of trees in different crown classes in the study sites (CD = Codominant, D = Dominant, E = Emergent, I = Intermediate, O = Overtopped).

**Figure 9:**
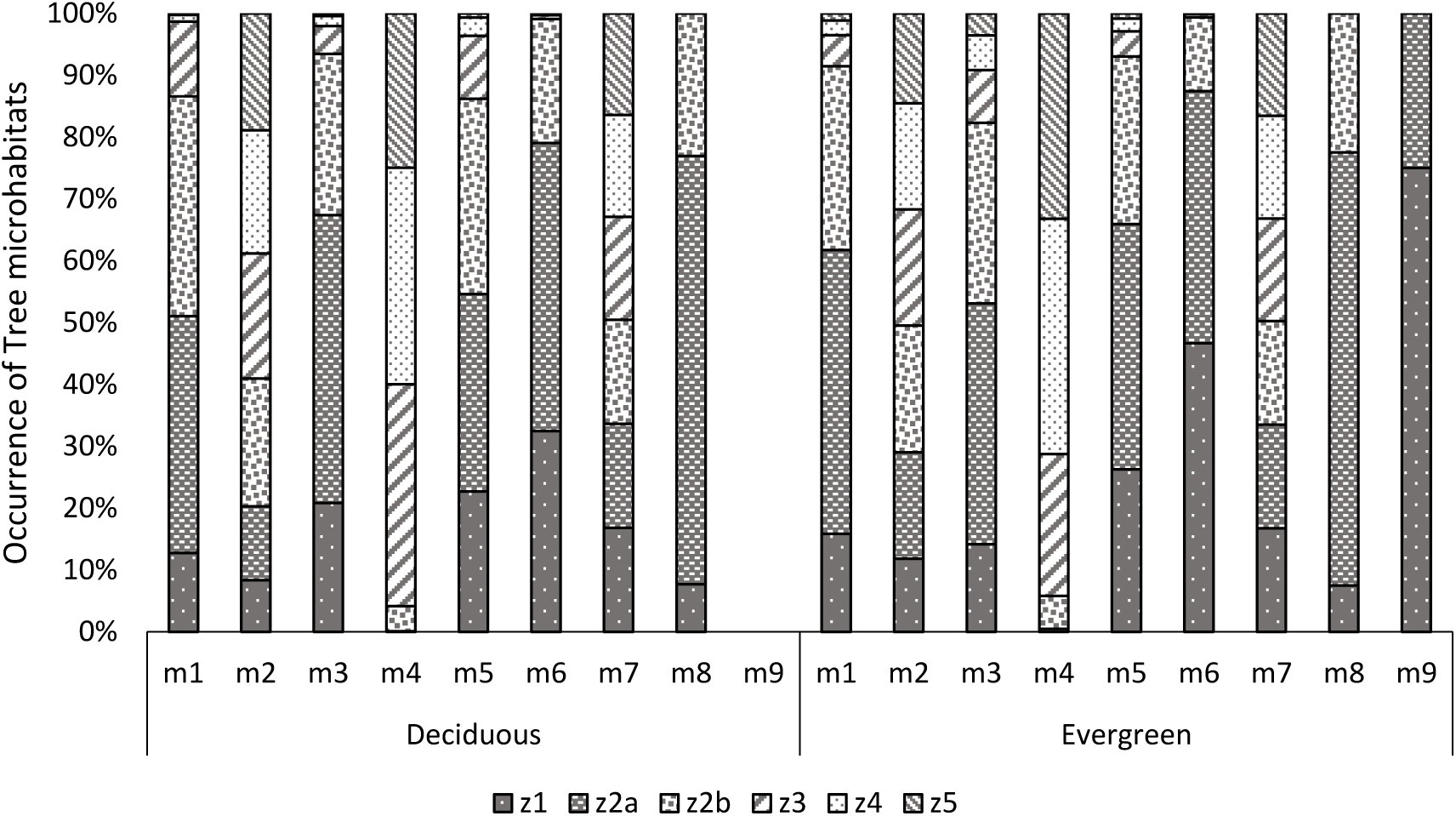
Vertical distribution of tree microhabitats (m1-m9 categories: see Table 2 for descriptions) in the studied habitats.

In deciduous habitat, 54.24 % of individuals were healthy, 39.61 % were unhealthy and 6.15 % were dead. Out of the sampled trees, 23.72 % of individuals were damaged by termites, 18.95 % were damaged by fungal decay and 57.33 % were damaged by other causes. Among the TMH host individuals, 24.12 % were damaged by termites, 18.93 % were decayed by fungi and 56.95 % were damaged by other causes. 13.72 % of individuals hosted cavities out of which 25.68% were damaged by termites, 26.23 % were having fungal decay and 48.09 % were damaged by other causes. The majority of the sampled trees were codominant (27.03 %), followed by intermediate (22.70 %), dominant and overtopped (each 20.00 %) and emergent (10.27 %). 25.99 % of TMH hosts were dominant, 22.34 % were overtopped, 21.05 % were codominant, 19.01 % were intermediate and 11.60 % were emergent. 25.69 % of cavity hosts were codominant, 21.56 % dominant, 21.10 % intermediate, 18.81 % overtopped and 12.84 % emergent (Figure 8). The difference in the dbh of TMH host and cavity host trees in various crown classes was highly significant (*F* = 195.258; *p*<0.001 and *F* = 32.208; *p* = <0.001 respectively), but post hoc analysis revealed that the difference in the dbh of trees in Intermediate and Codominant classes was not significant (*p* = 0.0753). The height of TMH host and cavity host trees differ significantly across the crown classes (*F* = 1069.177; *p*<0.001 and *F* = 203.662; *p*<0.001, respectively). Among the TMH host and cavity host trees, there was a significant difference between the basal area of trees in various crown classes (*F* = 115.7755; *p*<0.001 and *F* = 20.600; *p*<0.001, respectively). However, the post hoc analysis revealed that the basal area of Codominant cavity hosts did not differ significantly from individuals in the Intermediate class (*p* = 0.4899) and Overtopped cavity hosts did not differ significantly from the hosts in the Intermediate class (*p* = 0.4713). In deciduous, the total abundance of TMHs observed was 5,196 and epiphytic and epixylic structures were the most abundant type (1,306) with the highest abundance in the upper part of the trunk (Zone 2b) (270), followed by primary branches (Zone 3) (264), followed by Secondary branches (Zone 4) (260) (Figure 9).

## 4. Discussion

### 4.1. Diversity and abundance of tree microhabitats – the importance of host availability

All the tree microhabitat types reported in the temperate forests have been also observed during this inventory in tropical forests. Large sized trees are more likely to host tree microhabitats in any forest (Vuidot et al., 2011). The observations of this study in the tropical habitats were also consistent with this view. In tropical evergreen habitats, the average basal area of trees and TMH host trees estimated to be 67.03 sq m ha^-1^ and 54.31 sq m ha^-1^, respectively, in the present study were higher compared to the reports from few evergreen habitats of the Western Ghats 29-42 sq m ha^-1^ (Pascal and Pélissier, 1996), 39.7 sq m ha^-1^ (Pascal and Pélissier, 1996), 32.13 sq m ha^-1^ (Bhat et al., 2000), 43 sq m ha^-1^ (Rai, 2016). However, present estimations were lower compared to the basal area previously reported for SVNP 102.7 sq m ha^-1^ (Singh et al., 1984) and other evergreen habitats in the Western Ghats 94.64 sq m ha^-1^ (Parthasarathy, 1999) and 81.38 sq m ha^-1^ (Sundarapandian and Swamy, 2000). The average basal area estimated for trees in the moist deciduous habitat 37.77 sq m ha^-1^ was higher compared to previous reports from other moist deciduous habitats of the Western Ghats, 32.62 sq m ha^-1^ (Bhat et al., 2000), 28.05-33.77 sq m ha^-1^ (Sundarapandian and Swamy, 2000), 30-35 sq m ha^-1^ (Rai, 2016), 13.05-28.42 sq m ha^-1^ (Naidu et al., 2018). However, the basal area of TMH host trees in deciduous habitats were within the reported range 29.92 sq m ha^-1^. Probably the range of host trees for the tropics may not be limited to the range observed during this study, and more sampling in other sites may add more information on the relationship of host tree girth and TMH and cavity occurrence.

Majority of the previous studies compare the pattern of TMHs and its hosts in natural and managed stands to derive management interventions (Winter and Möller, 2008; Vuidot et al., 2011; Larrieu et al., 2014; Asbeck et al., 2021; Courbaud et al., 2022). Present study focused on the two major habitats in tropical region – evergreen and deciduous to develop a baseline information on the types of TMHs occurring in the tropics and assess their diversity. Based on previous studies it was hypothesised that evergreen stands by virtue of more density and high species richness will have higher diversity and abundances of TMHs. In terms of host species composition, evergreen habitats have more richness and diversity of species compared to the deciduous habitats. This is one of the major reasons for more abundance of TMHs in the evergreen habitats similar to the reports from temperate forests (Vuidot et al., 2011; Ouin et al., 2015). Also the abundance of unhealthy trees which are major hosts of tree microhabitats is high in evergreen which further increases the TMH density and abundance as observed by Paillet et al. (2018) in the strict reserves of lowland and mountain forests in France. Abiotic factors such as altitude, slope, presence of rocky outcrops and waterbodies influence the species composition and result in heterogeneity in the stands. This may influence the probability of occurrence of injuries, epiphytes and growth forms (Stokes et al., 2005; Dorren and Berger, 2006; Oliva and Colinas, 2010; Asbeck et al., 2022), as suggested in the ‘habitat heterogeneity hypothesis’ (MacArthur and MacArthur, 1961). Variations in such factors in the examined habitats may be resulted in alterations in the structure and composition of species and, consequently, the availability of suitable host tree species. The influence of host availability is more prominent in deciduous stands due to low relative abundance of species compared to the evergreen stands. In the studied habitats, nearly 50% of the species are hosting TMHs and cavities. Most of the tropical tree species tend to have aggregate distribution pattern (Condit et al., 2000). The diversity indices and distribution pattern indicate that some of the TMH sub-categories may have highly clustered distribution (e.g. epiphytes, cavities), and thus not recorded in all the quadrat and/or transects reducing the microhabitat diversity in both the habitats. If the TMHs have strong preference for the host species, then host and the TMH will follow similar abundance and distribution pattern, a reason for varying presence and abundances in the samples. Buttresses, flutes and similar growth forms support trees to withstand the wet soil conditions (Richter, 2014) and are characteristic of species of wet evergreen forests. In present study, such structures were one of the dominant TMHs in the evergreen habitat, but poorly represented in the deciduous stands.

### 4.2. The ‘Habitat Effect’ – horizontal and vertical distribution patterns

Between the evergreen and deciduous habitats, evergreen appear to have more naturalness due to its species richness and stand density and as expected it had higher TMH density compared to deciduous habitat. The higher density of TMHs per unit area in evergreen habitat compared to deciduous habitat, may be due to high abundance of host trees, availability of host in suitable vitality classes and diameter of trees (Larrieu and Cabanettes, 2012). In evergreen habitat, the density of TMHs per tree was 12.4±4.9 per tree while in deciduous it was 20.4±7.2 per tree. The stem density of host and non-host trees in evergreen habitats was higher compared to the deciduous which reduced the overall density of TMHs. However, the density of TMHs among the TMH host trees in the evergreen habitats was 2 per tree and 3 per tree in deciduous habitat. Probably host species are more important factor influencing the diversity and abundance of epiphytes such as lichens (Roper, 2018). But, the higher abundance of TMHs per tree in deciduous habitat may be due to proportionately higher abundance of individuals with poor vigour which make it prone to injuries and damages increasing the abundance of TMHs as observed in this study. Even the cavity excavators prefer species prone to fungal decay due to natural ageing or vulnerabilities (Cadieux et al., 2023). Availability of such factors increase the occurrence of TMHs such as cavities in a habitat.

With increase in the species richness in the stand the species richness of TMH host trees also increased. The richness of cavity host trees followed the stand species richness in deciduous, but in evergreen the cavity host pattern was different indicating host specificity as an important factor for occurrence of cavities. The diversity estimates for cavity host remained less than the stand diversity estimates probably because all present species may not host cavities and/or the number of individuals of host species may be less. Also, the number and abundance of host species differ in the two habitats; the TMH host may not necessarily host cavity. Species such as *Alseodaphne semecarpifolia*, *Cryptocarya bourdeillonii*, *Tabernaemontana heyneana* in evergreen habitat and *Morinda pubescens* and *Terminalia arjuna* in deciduous habitat do not host TMHs. Across the transects, in both the habitats there was high heterogeneity in the samples (Figure 7), similar pattern was observed for the host communities of TMH and cavities. While the samples represent uneven distribution of species and TMH host, the cavity host species show drop in evenness in both the habitats for the transects where the number of host species were less. The higher values of Berger-Parker index indicate that probably majority of the TMH and cavity hosts are species which are dominant in that habitat. Due to high singletons and/or doubletons, the Chao-1 estimate for the stand has high values in some of the transects. These rare species may not be hosting the TMHs which reduces the estimate value for host communities. The cavity host species are having even lower Chao-1 estimate compared to the TMH host indicating that probably the most abundant and dominant species are hosting cavities. The statistical differences in the diversity estimates for the stand and TMH and cavity host community were highly significant in both the habitats.

Within evergreen stands, plots and transects with higher diversity of species may have higher diversity and abundances of TMHs (Figure 7). In nature, stands with more abundance of host stems have more potential to provide hosts (Vuidot et al., 2011; Asbeck et al., 2022). On comparing the occurrences of trees, TMH and cavity host trees and TMHs including cavities it was observed that some of the tree microhabitats are abundant and more frequent irrespective of low tree and host abundances. Such as, in evergreen habitat the transect with lowest stand density (68.60 per plot) had highest density of TMHs host (60.40 per plot) and highest TMH density (162.20 per plot). But the same plot had lowest density of cavity hosts (approximately 4 per plot) and cavity density of 6 per plot compared to the highest density 9 per plot recorded in the habitat. It is probably due to the presence or absence of large trees or wood characteristics or suitable host species for cavities may not be present. This indicates the importance of stand composition (Vuidot et al., 2011), influence of host availability (Großmann et al., 2018), characteristics of potential hosts (Larrieu et al., 2012) and stand features related to naturalness for availability of suitable host for cavities as concluded by Winter and Moller (2008) and Asbeck et al. (2022).

All these stand parameters for the evergreen and deciduous habitats were found to vary significantly (Table 6). The estimated density of TMHs in tropical forest stands were higher 973 ha^-1^ (in evergreen habitat) and 767 ha^-1^ (combining both the habitats) than those reported in the natural stands of temperate forests by Michel and Winter (2009) (745 ha^-1^) and Ouin et al. (2015) (4.67 ± 0.78 per 100 m^2^ area). Michel and Winter (2009) identified a set of TMHs, such as broken tree top, bayonet top, crack/scars, bark loss, hollow tree trunk (rot holes), bark pockets, burl, heavy resinosis, bark burst as indicator TMHs of natural forest. In present study also TMH structures related to injuries and exposed wood had highest occurrence and tree growth forms made highest occurrence in the evergreen and might be an indicator of low disturbance in these stands. Tree growth forms had very low occurrence in deciduous stands, even in evergreen some of the specie tree species such as *Cullenia exarillata* and *Drypetes venusta* were frequently encountered with such structures as well as other TMHs such as cavities, dendrothelms, epiphytes and crown deadwood. Since these structures are species specific, increase in the abundance of such species may increase the abundance of TMHs associated with it. Asbeck et al. (2022) also reported that some of the species tend to host a diversity of TMHs increasing the TMH counts in the area. The abundance of deadwood (snags and logs) has been also identified as an indicator of stand age (Larrieu et al., 2014), however it is difficult to draw such inference from this study as sampling was done in an evergreen forest community which is at its climax stage. But the high density of snags in the evergreen stands compared to deciduous stands may be one of the reasons for increasing the occurrences of other TMHs such as cavities as observed by Paillet et al. (2018). Our estimates for tree cavities and its host are much less compared to the observations from tropical rainforests of China (Liu et al., 2019). Even though the numbers of cavity host trees were higher in the habitat studied by Liu et al. (2019), the proportion of total number of individuals contributing as cavity host in present study was higher. In present study, the proportion of individuals hosting cavities was 7.36 % in evergreen and 13.72 % in deciduous, while Liu et al. (2019) estimated it as 6.2 %. The reason for high estimates of tree cavities by Liu et al. (2019) may be due to sampling difference, as they documented cavities with a minimum entrance dimension ≥ 2 cm while we sampled cavities with minimum entrance dimension of 5 cm only.

Observation in this study shows that in both evergreen and deciduous habitats, trees with larger diameter and height classes are with an unhealthy vigour. The dead trees were more abundant in lower diameter and height classes or maybe they were broken by wind, storm, diseases, etc. Among the TMH host trees, in evergreen habitats the trees in larger diameter and height classes hosted TMHs, but in deciduous trees in even small diameter classes hosted TMHs (Figure 9). In deciduous habitat, stand density and diameter of trees are usually lesser than the evergreen habitats, also some of the tree species are more prone to have persistent injuries and damages (Romeiro et al., 2021) which may be one of the reasons for high number of TMH hosts even in lower diameter classes. Kõrkjas et al. (2021) also suggested that the lack of ability to heal the injuries make trees more potential to host.

Some of the TMHs facilitate microclimate and substrate conditions (Regnery et al., 2013), such as forked branches and rot holes for strangler figs and decay stage of snags for cavity excavation for formation of other TMHs as observed during this study and resource availability for the biota (Ganault et al., 2021) which might play key role in the ecosystem. The high proportion of TMH and cavity hosts in lower strata of evergreen habitat may be due to the effect of canopy induced microclimate (Angelini and Silliman, 2014) which increases the epiphyte occurrences and injuries due to falling of neighbouring trees and/or its branches on the trees beneath. The occurrences of some of the TMHs such as epiphytic and epixylic structures and Crematogaster ant nests are also governed by the microclimate features of the forests (Ramachandra et al., 2012; Elias et al., 2024) thereby influencing the diversity of tree microhabitats and their abundances in the habitats.

The results indicate that the presence of TMH and their relationship with the stand heterogeneities in respective habitats is not a random event, but there are some underlying factors resulting in this pattern. Further adding the view that the characteristics of the habitat including abiotic factors play an important role in the formation and occurrence of TMHs (Larrieu et al., 2012; Regnery et al., 2013; Asbeck et al., 2019; Augustynczik et al., 2019).

## 5. Conclusion

This study provides baseline information related to TMHs in the tropical forests. TMHs were observed in representative tropical evergreen and moist deciduous habitats in the southern Western Ghats region, grouped into 9 broad categories and 33 sub-categories. Evergreen forests had 9 TMH types with an average diversity of 79.31 %, including excrescences, growth forms, and epiphytic structures. These TMHs often depend on specific host species. Tree growth forms like flutes and buttresses can create concavities for microsoil accumulation, epiphytes, and nesting substrates. In deciduous forests, 8 TMH types were recorded (Crematogaster ant nests were absent) with 74.44 % diversity. Injuries, exposed wood, and epiphytic structures were most common. Injuries and exposed wood can facilitate other TMHs like insect galleries and fungal decay, increasing diversity. There were significant differences in tree density, TMH host trees, cavities, and other factors between the two habitats.

Evergreen had high stand basal area, snags, TMH host trees, TMHs per hectare, and cavities per hectare. Density of TMHs per tree was higher in deciduous, while cavity density per tree was similar in both habitats. Differences in stand characteristics like diameter, height, and basal area may explain higher TMH occurrences in evergreen. TMH counts were high in healthy trees in evergreen but high in unhealthy trees in deciduous. Tree strata and location of TMHs varied between habitats. This habitat had TMH host trees belonging to 105 species and 50 families, and 74 species and 35 families of cavity host trees. Deciduous habitat had 70 species and 35 families of TMH host trees and 44 species and 27 families of cavity host trees. TMH and cavity host trees were also the most abundant and dominant species in both habitats. Diversity indices of TMH host trees were high in both habitats, indicating diverse and abundant hosting species. Species richness of cavity hosts was about half that of TMH hosts, but frequency of TMH and cavity hosts was higher in deciduous.

During the study period the study area had two consecutive floods in 2018 and 2019 which interrupted the inventory and might have influenced the TMH occurrences due to falling of branches, trees, detachment of epiphytes etc. The tree microhabitats potentially related to the keystone structure ‘tree cavities’ were recorded. To avoid observer bias and reduce measurement error throughout the study period all the observations related to TMHs were done by one observer and measurement of tree and cavity measurements were done by one observer. However, this study is limited for the observations of TMHs in canopy such as cavities and epiphytes (attached to upper part of large branches) which were difficult to observe or confirm its presence from ground with the help of binocular. For these reasons, the TMH occurrences and abundance in upper zones may appear less compared to the lower zones contrary to the knowledge that canopies in tropical forests harbour more biological diversity.

In conclusion, TMH occurrence patterns in the tropics resemble other regions, but density and abundance are higher. Further investigations of tree microhabitats in different habitat types, the cooccurrences/correlated distribution patterns of TMHs in the tropical forests are needed to understand the structure and roles of such microhabitats for the conservation and management of tropical forests.

## CRediT authorship contribution statement

**Bharati Patel:** Conceptualization, Investigation, Data curation, Formal analysis, Methodology, Writing – original draft, Writing – review & editing. **Sreejith Sivaraman:** Investigation, Data curation, Methodology, Writing – review & editing. **Hrideek T.K.:** Writing – review & editing. **Peroth Balakrishnan:** Conceptualization, Investigation, Data curation, Formal analysis, Methodology, Funding acquisition, Writing – original draft, Writing – review & editing.

## Acknowledgment

We acknowledge financial support from the Ministry of Environment, Forests, and Climate Change, Govt. of India (Grant No. 14/34/2014-ERS/RE dated 02/06/2016), and thank the Kerala Forest and Wildlife Department for permissions and support.

