## Supplemental Appendix A for "Diversity and abundance of tree microhabitats in the tropical forests of southern Western Ghats, India"

### Supplementary material

#### APPENDIX A: Causes of un-healthiness of host trees other than termites and fugal decay identified during the study

| S.N. | Cause | S.N. | Cause | S.N. | Cause | S.N. | Cause |
|---|---|---|---|---|---|---|---|
| 1 | Bark damaged by natural growth of trees | 20 | Burls on stem | 39 | Dead branches on stem | 58 | Fire damage on stem |
| 2 | Bark pealed by animals or humans | 21 | Cankers on stem | 40 | Dead branches in canopy | 59 | Woodpecker foraging pecks on stem |
| 3 | Bark spit due to natural growth of trees or cankers exposing sapwood to infections | 22 | Burls and Canker on stem | 41 | Decay at base of the stem | 60 | Hollow stem, heart rot |
| 4 | Bark and sapwood both damaged by biotic or abiotic agents | 23 | Canker and bark split on stem | 42 | Fissure on stem | 61 | Host dead and strangler Ficus spp. Covered the tree |
| 5 | Bark peal with splits on stem | 24 | Canker damage in bark | 43 | Decay and fissures on stem | 62 | Insect boreholes |
| 6 | Bark peal and damage to sapwood | 25 | Canker rot on stem | 44 | Fissures and rot hole on stem | 63 | Insect bore holes with saprun on stem |
| 7 | Bark pealed and decay in sapwood | 26 | Cankers and branch knots on stem | 45 | Decay in bark and sapwood at branch knot | 64 | Liana completely covered the tree |
| 8 | Bark damaged and decay in sapwood | 27 | Canopy branches dead | 46 | Decay in branch knots | 65 | Saprun from stem |
| 9 | Bark and sapwood damaged and decay | 28 | Canopy damage by liana | 47 | Decay in buttress | 66 | Rot holes on stem |
| 10 | Bark damaged by collision with neighbouring tree | 29 | Canopy partly dead | 48 | Decay in forking due to rainwater | 67 | Rot holes and saprun on stem |
| 11 | Bark and stem damage or decay by fallen neighbouring tree | 30 | Cavities on stem due to heartwood decay | 49 | Decay in heartwood | 68 | Saprun from rotten heartwood of stem |
| 12 | Bark damage and decay in sapwood | 31 | Cavity formed by decay in buttress | 50 | Crooked stem | 69 | Stem hollow by termite decay |
| 13 | Bark damage and saprun | 32 | Cavity formed by decay in stem | 51 | Denderothelm cavity in stem, decay | 70 | Termite and burls on stem |
| 14 | Bark splits with saprun | 33 | Cavity formed by decay on stem filled with water | 52 | Denderothelm in broken branch | 71 | Tree top broken, splintered stem |
| 15 | Bark damage due to rot | 34 | Covered with strangler Ficus | 53 | Denderothelm in dead branches | 72 | Stem hollow, cavity excavated to collect honey from hollow stem |
| 16 | Bark damaged and dead branches on stem | 35 | Dead branch knots on stem | 54 | Epiphyte Ficus covers the host tree | 73 | Unknown |
| 17 | Bark/stem damaged by liana | 36 | Dead branches and decay cavity | 55 | Excavated cavities on stem | 74 | Decay due to rainwater |
| 18 | Branches dead due to impact of fallen neighbour tree | 37 | Dead branches and epiphytes in canopy | 56 | Exposed heartwood | 75 | Splintered stem |
| 19 | Bulges on stem | 38 | Dead branches and rot holes in stem | 57 | Stem decay in heartwood | 76 | Axe-marks on stem |
