## Supplemental Appendix B for "Diversity and abundance of tree microhabitats in the tropical forests of southern Western Ghats, India"

### APPENDIX B: Methods used for the estimation of stand characteristics

a) Stand basal area (G) ($m^2ha^{-1}$)

Basal area is usually calculated from the diameter (m) calculated from girth measured at breast height (1.37 m) of the trees. In this study the diameter of the trees was in meters and area of the plot (sample) was 1 ha (10000 sq m) the stand basal area was calculated by adding the basal area of all the trees in a transect.

b) Stand Diameter (D) (m)

The diameter of trees is measured from girth measured at breast height (1.37 m) of the trees. Since, the diameter of the trees was not showing normal distribution, to avoid bias due to large variations the diameter of the tree was weighted by its basal area. For this Lorey's formula was applied to calculate *d* which is multiplication of diameter of tree at breast height of the individual tree and its basal area which was divided by the total basal area of the stand (Lorey 1878; dos Santos Vieira et al. 2020). In this study the diameter of the trees was in meters and area of the plot (sample) was 1 ha (10000 sq m) the stand diameter was calculated by adding the diameter of all the trees in a transect calculated by using Lorey's formula.

c) Stand Height (H) (Lorey's Formula) (m)

The height of trees (m) is measured from ground level. Since, the height of the trees was not showing normal distribution, to avoid bias due to large variations the height of the tree was weighted by its basal area. For this Lorey's formula was applied to calculate *h* which is multiplication of height of tree and its basal area, and it was divided by the total basal area of the stand (Lorey 1878; dos Santos Vieira et al. 2020). In this study the height of the trees was in meters and area of the plot (sample) was 1 ha (10000 sq m) the stand height was calculated by adding the height of all the trees in a transect calculated by using Lorey's formula.

d) Stand Density (D)

It shows the degree of crowding of stem in a plot.

$$D = \frac{\sum_{i=1}^{n} n}{Area\ of\ the\ plot}$$

*n* = number of individuals in the stand

In this study the area of the plot (sample) was 1 ha (10000 sq m) the stand density was calculated with the number of trees in a transect
