## Supplemental Appendix C for "Diversity and abundance of tree microhabitats in the tropical forests of southern Western Ghats, India"

### APPENDIX C: Method used for estimation of diversity indices

Diversity indices were estimated using standard formulae for all the recorded trees, tree microhabitat host trees, cavity host trees and snags. For tree microhabitats (overall and category-wise) only frequency (%), density (counts ha$^{-1}$) and abundance were calculated at the units - per tree and per ha (Curtis and McIntosh 1950). The diversity estimates were done following Magurran (2003). Microhabitat Diversity Index was calculated for nine tree microhabitat categories as well as 33 sub-categories following Paillet et al. (2018). Following are the formulas used for this analysis:

**a)** Frequency

It measures the proportion of how frequently a species is encountered in the samples based on presence absence data.

Frequency % of tree microhabitat tree (per ha) = $\frac{Number\ of\ quadrats\ in\ which\ tmh\ tree\ occurred}{Total\ number\ of\ quadrats\ studied} * 100$

Frequency % of cavity tree (per ha) = $\frac{Number\ of\ quadrats\ in\ which\ cavity\ tree\ occurred}{Total\ number\ of\ quadrats\ studied} * 100$

Frequency % of dead tree (per ha) = $\frac{Number\ of\ quadrats\ in\ which\ dead\ tree\ occurred}{Total\ number\ of\ quadrats\ studied} * 100$

Frequency % of tree microhabitat (per tree) = $\frac{Number\ of\ trees\ on\ which\ the\ tmh\ occurred}{Total\ number\ of\ trees\ studied} * 100$

**b)** Density

Estimates the distribution of species per unit area across the samples based on count data.

Density of tree microhabitat tree (per ha) = $\frac{Total\ number\ of\ the\ tmh\ trees}{Total\ number\ of\ quadrats\ per\ unit\ studied}$

Relative Density of tree microhabitat trees (per ha) = $\frac{Total\ number\ of\ the\ tmh\ trees}{Total\ density\ of\ all\ the\ tmh\ trees} * 100$

Density of cavity tree (per ha) = $\frac{Total\ number\ of\ the\ cavity\ trees}{Total\ number\ of\ quadrats\ per\ unit\ studied}$

Relative Density of cavity tree (per ha) = $\frac{Total\ number\ of\ the\ cavity\ trees}{Total\ density\ of\ all\ the\ cavity\ trees} * 100$

Density of dead tree (per ha) = $\frac{Total\ number\ of\ the\ dead\ trees}{Total\ number\ of\ quadrats\ per\ unit\ studied}$

Relative Density of dead trees (per ha) = $\frac{Total\ number\ of\ the\ dead\ trees}{Total\ density\ of\ all\ the\ dead\ trees} * 100$

Density of tree microhabitat (per tree) = $\frac{Total\ number\ of\ the\ tmh}{Total\ number\ of\ tmhs\ per\ unit\ studied}$

Relative Density of tree microhabitat (per tree) = $\frac{Density\ of\ a\ given\ tmh}{Total\ density\ of\ all\ the\ tmhs} * 100$

Density of tree microhabitat (per ha) = $\frac{Total\ number\ of\ the\ tmh}{Total\ number\ of\ quadrats\ per\ unit\ studied}$

Relative Density of tree microhabitat (per ha) = $\frac{Total\ number\ of\ the\ tmh}{Total\ density\ of\ all\ the\ tmhs} * 100$

**c)** Abundance

It estimates the occurrence of a species per unit area based on count data and number of samples it was encountered.

Abundance of tree microhabitat tree (per ha) = $\frac{Total\ number\ of\ tmh\ trees}{Total\ number\ of\ quadrats\ per\ unit\ in\ which\ they\ occurred}$

Relative Abundance of tree microhabitat tree (per ha) = $\frac{Total\ number\ of\ tmh\ trees}{Total\ number\ of\ individuals\ of\ all\ the\ trees} * 100$

Abundance of cavity tree (per ha) = $\frac{Total\ number\ of\ cavity\ trees}{Total\ number\ of\ quadrats\ per\ unit\ in\ which\ they\ occurred}$

Relative Abundance of cavity tree (per ha) = $\frac{Total\ number\ of\ cavity\ trees}{Total\ number\ of\ individuals\ of\ all\ the\ trees} * 100$

Abundance of dead tree (per ha) = $\frac{Total\ number\ of\ dead\ trees}{Total\ number\ of\ quadrats\ per\ unit\ in\ which\ they\ occurred}$

Relative Abundance of dead tree (per ha) = $\frac{Total\ number\ of\ dead\ trees}{Total\ number\ of\ individuals\ of\ all\ the\ trees} * 100$

Abundance of tree microhabitat (per tree) = $\frac{Total\ number\ of\ the\ tmh\ recorded}{Total\ number\ of\ trees\ per\ unit\ on\ which\ they\ occurred}$

Relative Abundance of tree microhabitat (per tree) = $\frac{Total\ number\ of\ the\ tmh\ recorded}{Total\ number\ of\ all\ the\ tmhs\ on\ all\ the\ trees} * 100$

Abundance of tree microhabitat (per ha) = $\frac{Total\ number\ of\ the\ tmh}{Total\ number\ of\ quadrats\ per\ unit\ in\ which\ they\ occurred}$

Relative Abundance of tree microhabitat (per ha) = $\frac{Total\ number\ of\ the\ tmh}{Total\ number\ of\ all\ the\ tmhs\ in\ all\ the\ quadrats} * 100$

**d)** Species Richness

The number of taxa/species in a sample represent the species richness of the sample. The higher the value, higher is the number of species/taxa:

Number of taxa ($S$)

**e)** Dominance Index

The dominance index indicates it the sampling unit is dominated of one or a few species and it ranges from 0 to 1. Higher value of this index (close to 1) indicates infinite diversity and with value close to 0, the sample is expected to have low diversity.

Simpson Index (D) (Simpson 1949):

$D = 1 - \frac{\sum_i n_i(n_i-1)}{n(n-1)}$ $n_i$ = Number of individuals in the $i$-th species; $n$ = Total number of individuals in the community.

Dominance Index = 1−Simpson index

Inverse Simpson has its lowest value as 1, and higher the value of this index higher is the diversity.

**f)** Shannon Diversity Index (*H)* (Shannon and Wiener, 1963)

The index is based on the hypothesis that the sampling has been done randomly; the higher values of this index indicates that the sample has high diversity.

$$H = -\sum_{i} \frac{n_i}{n} \ln \frac{n_i}{n}$$

*ln* = natural log; $n_i$ = Number of individuals in the *i*-th species; *n* = Total number of individuals in the community

**g)** Equitability Index ($E_H$) (Sheldon, 1969)

It measures the evenness of species in the sampled community. It quantifies the similarities in the abundances of all the species in the community. The value ranges from 0 to 1, 0 being highly uneven state of species distribution and 1 being highly even distribution of species.

$\boldsymbol{E_H}$ = $H/\ln(S)$……………$H$ = Shannon diversity Index; $S$ = the number of taxa.

**h)** Jaccard Similarity Index (*J*) (Jaccard 1901, 1912)

The similarities in the occurrences and co-occurrences of species and families in the samples (transects) were calculated as a coefficient using the formula:

$$J = \frac{a}{a+b+c}$$

*a* = number of species common in both the samples, *b* = number of species unique to 1$^{st}$ sample, *c* = number of species unique to 2$^{nd}$ sample

**i)** Fisher's Alpha (Fisher et al. 1943)

It is a parametric index of diversity and is believed to be useful for even for small sample size. It establishes the relationship between the number of number of species and number of individuals of the species. It is found suitable for species which follow log series model and it is assumed that one or a few factors determine the properties of the community.

$S = a * ln\left(1 + \frac{n}{a}\right)$ *S* = number of taxa, *n* = number of individuals; *a* = Fisher's alpha

**j)** Berger-Parker Dominance Index (*d*) (Berger and Parker, 1970)

This index measures the numerical importance of most abundant species in the sample.

d = $N_{max}/N$

$N_{max}$ = number of individuals in the most abundant species; *N* = total number of individuals in the sample.

The higher value of this index indicates that the community is dominated by the most common species and there is no even distribution of species.

**k)** Chao1 (Chao, 1984), bias corrected and is an estimate of total species richness based on the numbers of singleton and doubleton species. If the sample contains more singletons, the index will estimate greater species richness compared to sample without rare species, indicating that it is likely that more undetected species exist.

$S_1 = S_{obs} + \frac{F_1^2}{2F_2}$ where, $S_{obs}$ = number of species in the sample; F1 = number of singletons and F2 = number of doubletons

All the biodiversity indices were calculated for stand and transect level for comparisons using PAST software (Hammer et al., 2001).

**l)** Microhabitat Diversity Index (%)

The diversity index for tree microhabitats were calculated for each plot and then averaged to per hectare (transect level) for comparison of stands. The microhabitat diversity index was calculated following Paillet et al. (2018):

$$di = \frac{Number\ of\ tree\ microhabitat\ types\ observed\ in\ the\ plot}{Maximum\ number\ of\ tree\ microhabitat\ types\ expected\ to\ be\ observed} \times 100$$

Based on observations, in this study for diversity index of categories (di1) the denominator was 9 and for diversity of index of sub-categories (di2) the denominator was 33.

**m)** Coverage Value Index (IVC)

The index gives numerical representation of the percentage coverage of species or community. It is calculated as follows (Mueller-Dombois and Ellenberg 1974):

IVC = Relative Density (RD) + Relative Dominance (RDo)

IVC % = (IVC/200) × 100

**n)** Importance Value Index (IVI)

The index value represents the important species in the area. It is calculated as follows (Mueller-Dombois and Ellemberg 1974):

IVI = Relative Frequency (RF) + Relative Density (RD) + Relative Dominance (RDo)

The value of IVI reaches up to 300

IVI % = (IVI/300) × 100
